# Regulation of plasma membrane tension through hydrostatic pressure and actin protrusion forces

**DOI:** 10.1101/2025.03.24.645030

**Authors:** Yogishree Arabinda Panda, Elisabeth Fischer-Friedrich

**Author notes:** Electronic mail: Corresponding author.

## Abstract

The plasma membrane and its associated proteins form a critical signaling hub, mediating communication between the extracellular environment and the intracellular space. Previous research suggests that both membrane trafficking and signaling activity are influenced by mechanical tension in the plasma membrane. Despite its importance, the mechanisms by which cells regulate membrane tension remain poorly understood. Using the optical tension sensor FliptR and AFM-assisted tether force measurements, we investigate plasma membrane tension regulation in mitotic cells by measuring tension changes following cytoskeletal and cell shape perturbations. Our findings show that in both assays, reported tensions are critically influenced by the cytoskeleton, however, with partially deviating trends highlighting the conceptual differences between bare and apparent membrane tension. By integrating experimental data with theoretical modeling, we demonstrate that the actin cytoskeleton regulates bare membrane tension through two distinct mechanisms: (i) modulation of intracellular hydrostatic pressure and (ii) adjustment of polymerization forces in actin-rich finger-like protrusions.

**SIGNIFICANCE STATEMENT:** The plasma membrane is a vital cell interface, facilitating communication with the environment. Mechanical tension in the membrane plays a crucial role in tuning this communication, yet its regulation remains poorly understood. This is partly due to the lack of precise measurement tools and the misleading use of membrane tether forces as proxies. In this study, we use advanced techniques — FliptR tension sensors, AFM-based tether force measurements, and theoretical modeling — to explore membrane tension in mitotically arrested cells. We find that the actin cytoskeleton regulates membrane tension via intracellular pressure control and polymerization forces in protrusions. Moreover, we show that tether forces do not reliably reflect in-plane membrane tension, making them unsuitable as direct reporters of membrane mechanics.

## I INTRODUCTION

The plasma membrane is a lipid bilayer that consists of phospholipids and cholesterol as well as transmembrane and membrane-bound proteins^1,2^. It serves as a diffusion barrier for ions and organic molecules and creates a dynamic platform for cellular signaling and interactions with the environment^1–3^. Therefore, the plasma membrane has a key role in the regulation of physiological processes such as differentiation, cell migration, and cell proliferation^4^.

The plasma membrane is mechanically supported by the neighboring actin cortex - a thin cross-linked layer of actin polymers which is mechanically attached to the plasma membrane by diverse molecular interactions including ERM proteins, integrin complexes, formins and Arp2/3 ^5–9^. Myosin motor proteins introduce active contractile tension in the actin cortex thereby regulating the hydrostatic pressure excess in the cell^10–12^. The actin cytoskeleton further plays an important role in regulating membrane reservoirs by tethering caveolae^13^ and scaffolding finger-like protrusions such as microvilli and filopodia^14^.

While the plasma membrane is a fluid-like sheet due to lateral mobility of lipids, it has a considerable area compressibility modulus of ≈ 300 mN/m featuring the membrane as essentially inextensible^15^. Recent evidence highlights that mechanical tension in the plasma membrane plays a crucial role in modulating membrane-mediated signaling^16^. Notable examples include the mechanosensitive ion channel PIEZO, a pivotal regulator of epithelial homeostasis^17,18^, and G-protein-coupled receptors (GPCRs), the largest family of human membrane proteins, which are essential for regulating vision, olfaction, and taste^16^.

Membrane tension in cells can currently be measured with two methods: (i) tension-sensitive, membrane-associated optical sensors^19,20^, and (ii) the mechanical pulling of membrane tethers with concomitant force measurements using optical tweezers or atomic force microscope cantilevers^21,22^. While the first method holds the potential to measure changes of actual (bare) in-plane tension *γ*_mem_, the latter only measures apparent membrane tension *γ*_app_ = *γ*_adh_ + *γ*_mem_, which includes additional contributions from cortex-membrane adhesion energy *γ*_adh_. In a recent work, De Belly *et al*. demonstrated that actin cortex contractility and actin protrusions, driven by the activation of RAC or RHOA, can significantly alter apparent membrane tension with quick propagation through the cell upon local perturbations^21^. In this study, we combine atomic force microscopy (AFM) with fluorescence lifetime imaging microscopy (FLIM) of the sensor FliptR to investigate changes in bare membrane tension, apparent membrane tension, cortical tension, and filopodial dynamics in response to osmotic volume changes, as well as perturbations of the cytoskeleton and cell-substrate contact area. Our findings not only elucidate the conceptual and factual distinctions between bare and apparent membrane tension but also present a biomechanical scheme that explains how membrane tension responds to modifications in actin-cytoskeletal tension, intracellular hydrostatic pressure, and protrusive actin polymerization.

## II RESULTS

### A. Osmotic shocks trigger significant changes in FliptR lifetime

To investigate bare membrane tension, we used the fluorescent membrane tension probe FliptR. Earlier work characterized fluorescence lifetime changes of this dye in response to bare membrane tension changes in cells^19,23–27^; this effect was accounted for by the dependence of FliptR fluorescence lifetime on the orientation between its chromophoric groups^19^. For brevity, we will refer in the following to ‘bare membrane tension’ as just ‘membrane tension’. To validate the sensitivity of FliptR fluorescence lifetime to membrane tension in our biological model system - HeLa cells in mitotic arrest - we applied osmotic shocks and measured concomitant changes in cellular volume, surface area, and FliptR fluorescence lifetime (see Fig. 1a-f and Materials and Methods). Hypo-osmotic swelling increases cell volume and surface area, which supposedly increases membrane tension. Conversely, hyper-osmotic shrinkage reduces cell surface area, likely leading to a decrease in membrane tension. Fig. 1h,i displays exemplary transmitted light pictures of cells before and after osmotic shock. As shown in Fig. 1d, the emergent cell volume 500 − 600 s after the shock is inversely proportional to the osmolarity of the surrounding medium. Concomitantly, analysing FliptR fluorescence in the outer cortex-associated plasma membrane of cells, we can observe an increase in FliptR lifetime upon cell swelling following hypo-osmotic shock, and conversely, a FliptR lifetime decrease during hyper-osmotic shock (Fig. 1e,f). Correspondingly, FliptR lifetime after the shock demonstrated a negative linear correlation with the osmolarity of the medium, see Fig. 1e. Furthermore, emergent FliptR lifetimes correlate positively with emergent cell surface areas, exhibiting an approximately linear relation within ± 40% area change from the control state of isotonic medium, see Fig. 1f. The time evolution of cell size and FliptR lifetime upon osmotic shock is depicted in Fig. 1g. (Shocks are applied at time point *t* = 0 s.) In conclusion, we observe that FliptR fluorescence lifetimes change in positive correlation with expected membrane tension changes upon osmotic shock. Furthermore, our measurements suggest that membrane tension changes are not only quickly translated into FliptR lifetime changes but that corresponding changes are also lasting long term, i.e. on time scales of hundreds of seconds, see Fig. 1g.

**Figure 1.**
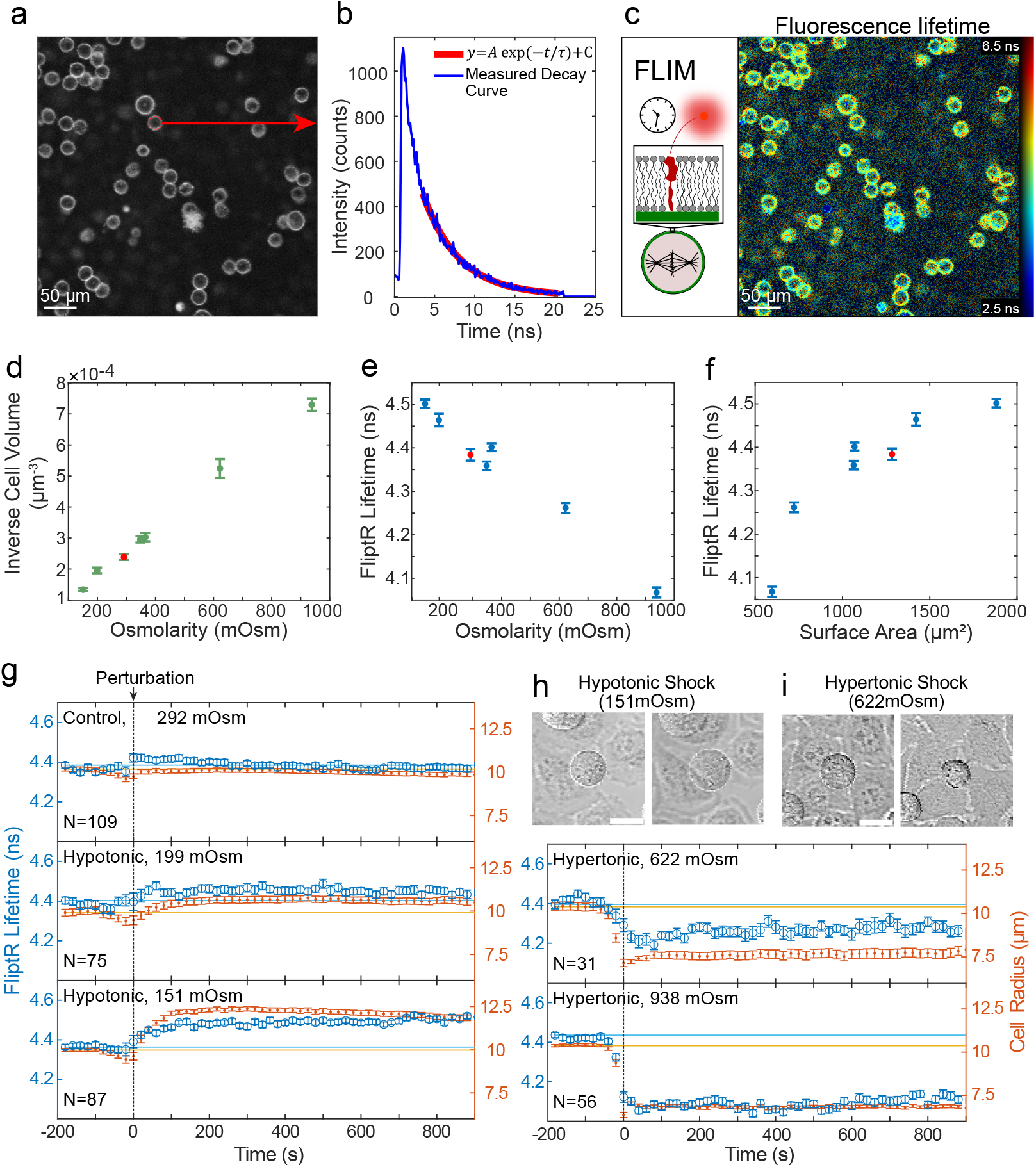
FliptR lifetimes change in correlation with cell surface area upon osmotic shocks. To prevent cells from progressing through the cell cycle, HeLa cells were in mitotic arrest. a) Confocal microscopy image of FliptR fluorescence in the equatorial plane of HeLa cells in mitotic arrest. The red circle shows the edge detection of the mitotic cell selected for analysis. Scale bar: 50 *µ*m. b) Histogram of photon detection times from the cortical region of the selected cell (blue curve). The mono-exponential tail fit (red curve) starts 2.5 ns after the peak and provides a fitted fluorescence lifetime, see Materials and Methods. c) Confocal image showing pixel-wise fitted FliptR lifetimes. Color intensity was scaled with pixel brightness such that regions with low photon counts appear dark. Scale bar: 50 *µ*m. d) Median inverse cell volumes after the osmotic shock scale approximately linearly with newly acquired osmolarities of the medium. Inverse volumes were averaged in the time interval 500 − 600 s after the shock. e) Median FliptR lifetimes from cell cortices scale approximately linear with newly established medium osmolarities. FliptR lifetimes were averaged in the time interval 500 − 600 s after the shock. f) Median FliptR lifetimes from cell cortices change in correlation with newly acquired projected cortex area. FliptR lifetimes and cortex areas were averaged in the time interval 500 − 600 s after the shock. d-f) Red data points indicate the control measurement where pure, isotonic medium was added to the dish during the experiment. g) Time evolution of cortical FliptR lifetime and cell radius during osmotic shock experiments. During control experiments, medium of the same osmolarity was added to the dish. Solid blue and orange lines indicate the average lifetime and cell radius before the perturbation. d-g) Each cell measured gave rise to one data point of the statistics. Numbers of measured cells are shown in panel g. Error bars show the standard error of the median. h,i) Transmitted light pictures showing a mitotic HeLa cell before and 500 s after an osmotic shock of indicated magnitude. Scale bars: 20 *µ*m.

### B. Impact of cytoskeletal inhibitors on FliptR lifetimes in the plasma membrane

In order to investigate the impact of the actin cytoskeleton on plasma membrane tension, we treated HeLa cells in mitotic arrest with various cytoskeletal drugs to selectively disrupt different components of the cytoskeletal network in the presence of the membrane tension sensor FliptR. Drugs were added to the medium after ≈ 200 s measurement time, i.e. at time point *t* ≈ 0 s. We then analysed the changes of FliptR fluorescence lifetime upon adding the drug to the medium. FliptR lifetime analysis was restricted to the fluorescence signal from the outer rim of the cell, i.e. the signal from the cortex-associated plasma membrane, see Materials and Methods. As a control, we added just medium without drugs to the well, see Fig. 2, top left panel. For control measurements, we observe a short-term increase in FliptR lifetime within the 0 − 200 s interval, likely caused by medium fluxes and the resulting increase in membrane tension. Next, we tested the influence of myosin II, which is the main generator of actin cytoskeleton contractility and cortical tension^28^. Upon addition of the myosin inhibitor Para-Nitro Blebbistatin (PN-Blebbistatin, 10 *µ*M), we find a persistent reduction of FliptR lifetime indicating a reduction of membrane tension (Fig. 2, blue curve). By contrast, inhibition of ROCK with Y27632 (25 *µ*M) did not induce changes in FliptR lifetime that were distinct from the control measurement (Fig. 2, dark red curve). Perturbing actin polymerization through inhibition of actin nucleators formin or Arp2/3 with their respective pharmacological inhibitors SMIFH2 (40 *µ*M) and CK666 (50 *µ*M) reduced FliptR lifetime suggesting a reduction in membrane tension (Fig. 2, violet and green curves). Further, we tested the influence of Ezrin- Radixin-Moesin (ERM) using the ERM inhibitor NSC668394^29^. Adding NSC668394 (50 *µ*M) to the cell medium, we find also a reduction of FliptR lifetime indicative of concomitant membrane tension decrease (Fig. 2, orange curve). Lastly, we inhibited microtubule polymerization using Nocodazole (5 *µ*M) and observed that Nocodazole-induced microtubule depolymerization caused a slight decrease in FliptR lifetime, indicating a corresponding reduction in membrane tension (Fig. 2, yellow curve). In conclusion, these results demonstrate a critical role of the actin and microtubule cytoskeleton in regulating plasma membrane tension.

**Figure 2.**
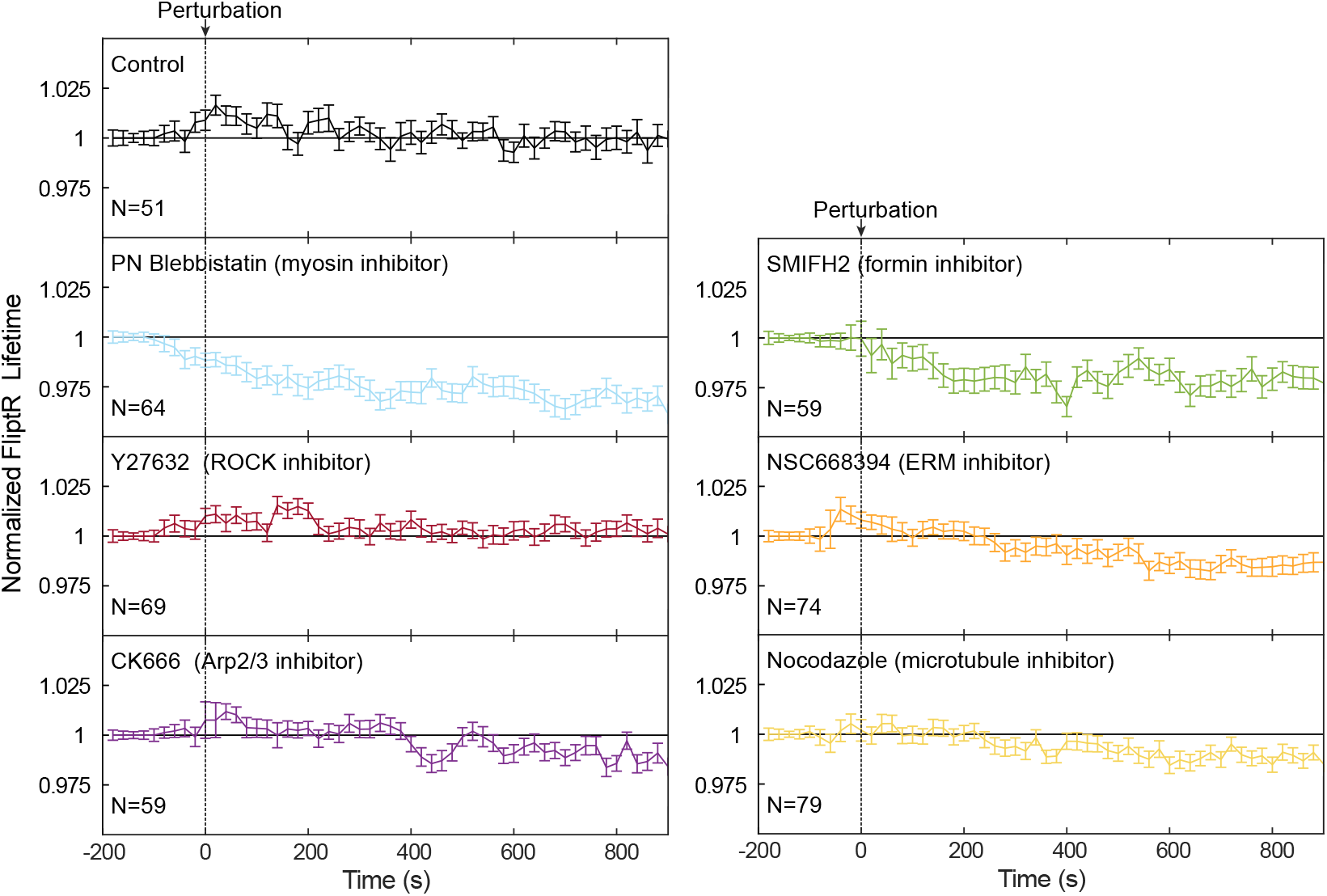
Time evolution of FliptR lifetimes in the plasma membrane of HeLa cells upon addition of cytoskeletal drugs. Cytoskeletal drugs were added after 200 s of measurement time, i.e. at time point *t* = 0 s. During control experiments (top left panel), pure cell culture medium was added at time point 0 s. To reduce the impact of cell-cell variations, FliptR lifetimes were normalized by their initial value (median of the first 8 time points). Images were recorded at a time interval of 20 s, see Materials and Methods. Error bars show the standard error of the median. Concentrations of cytoskeletal drugs used were as follows: Para-Nitro Blebbistatin (PN-Blebbistatin, 10 *µ*M), Y27632 (25 *µ*M), CK666 (50 *µ*M), SMIFH2 (40 *µ*M), NSC668394 (50 *µ*M), and Nocodazole (5 *µ*M).

### C. Increasing cell-substrate contact using uniaxial AFM compression leads to a decrease in FliptR lifetimes in the plasma membrane

Beyond the effects of osmotic shocks and cytoskeletal perturbations, we explored how membrane tension is influenced by cell shape and cell-substrate contact. To investigate this, we deformed HeLa cells in mitotic arrest using the cantilever of an atomic force microscope (AFM), see Fig. 3a. HeLa cells in mitotic arrest were initially confined between a glass cover slide and a wedged cantilever. By lowering the cantilever, we induced uniaxial compression and simultaneously measured the AFM force as a reporter of cortical tension, see Fig. 3b, Materials and Methods and^30,31^. To avoid blebbing during this process, cells were treated with the ROCK-inhibitor Y27632 leading to a reduced cortical tension. Previous studies showed that this uniaxial cell compression leads to cellular surface area increase while leaving cellular volume approximately constant^11,32^. As was shown before, cell shapes are well captured by shapes of minimal surface area and vanishing adhesion^11^. Therefore, theory predicts a total cell surface area increase of ≈ 2% upon uniaxial compression for a typical cell radius of ≈ 10 *µ*m and height changes as depicted in Fig. 3a. At the same time, however, the free-standing unsupported cell surface area is predicted to decrease by ≈ 3.5% as more membrane is included in the enlarged contact area upon compression. During this experiment, we measured FliptR lifetimes as a reporter of membrane tension through continuous FLIM imaging, see Fig. 3c and Materials and Methods. Despite large fluctuations of FliptR lifetimes and significant cell-cell variations, we find that on average

**Figure 3.**
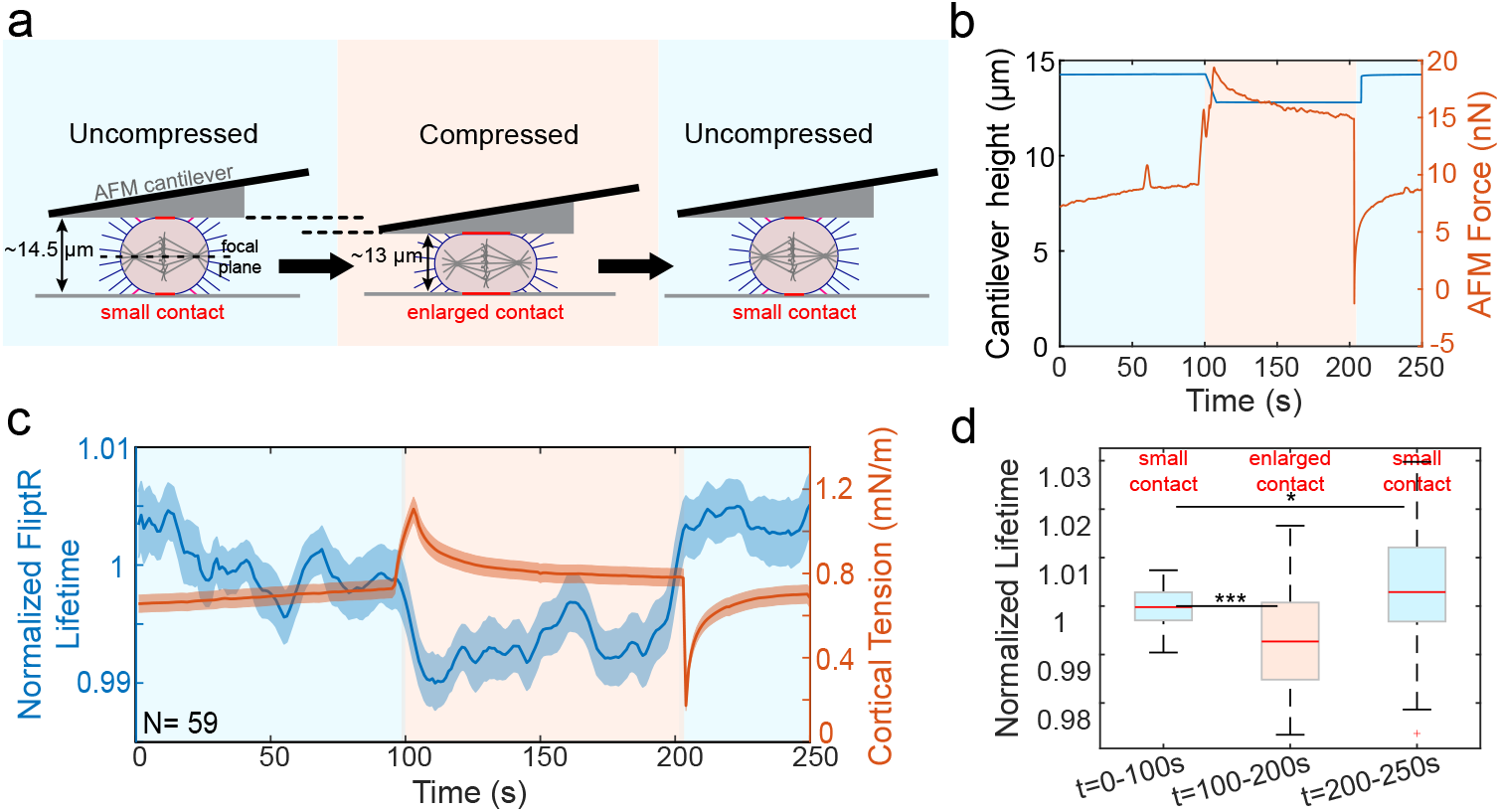
Membrane tension in the plasma membrane decreases in response to uniaxial cell compression and concomitant contact area increase. a) Schematic of the cell confinement assay. A HeLa cell in mitotic arrest is deformed with an AFM cantilever with a step of uniaxial compression, see Materials and Methods. In this way, the surface area of the cell is elevated, and cortical tension is temporarily increased. The free-standing membrane surface is reduced in this process and filopodia close to the periphery of the initial contact area (labelled in pink) are integrated into the contact interface in the compressed state. To prevent cell-blebbing during the process, the ROCK inhibitor Y-27632 (10 *µ*M) has been added to the medium. b) AFM force and cantilever height change over time for an exemplary cell. c) Population averages of cortical tension and normalized FliptR lifetimes over time for 59 cells. d) Boxplots of time-averaged normalized FliptR lifetimes for 3 time intervals (see panel c): before cell deformation (0 − 100 s), during cell deformation (100 − 200 s) and after cell deformation (200 − 250 s). We find that FliptR lifetimes are lower in the more strongly confined state and slightly elevated right after cantilever lifting. Significance was tested using a two-tailed Mann-Whitney U test and *p*≤ 0.05.

FliptR lifetimes decrease as cells are uniaxially compressed (*t* = 100 s) followed by the restoration and slight increase of lifetimes upon release of the compression (*t* = 200 s), see Fig. 3c,d. This observation suggests a membrane tension decrease upon cell compression and an increase upon its release. In particular, the deformation-induced peak in cortex tension right after cortex area increase upon lowering of the cantilever (see orange curve at *t* = 100 − 120 s) was not reflected in a correlated change of FliptR lifetime (see orange curve at *t* = 100 − 150 s), see Fig. 3c. We propose that a compression-induced decrease in membrane tension is triggered by a loss of filopodia at the rim of the initially free-standing cell surface area, which becomes part of the substrate contact upon compression. This localized loss of filopodia effectively reduces filopodia number, leading to a corresponding decrease in actin polymerization pressure and, consequently, lower membrane tension (see Section III for a theoretical reconsideration of this aspect). We verified the loss of filopodia in newly established contact areas through confocal imaging of Lifeact-mScarlet at the upper surface of the cell, see Fig. S2.

### D. Impact of cytoskeletal inhibitors on membrane tether forces

Previous research has presented the measurement of membrane tether forces as a reporter of apparent membrane tension 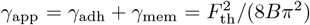, where *γ*_adh_ is the membrane-cortex adhesion energy per unit area, *γ*_mem_ is the in-plane membrane tension, *B* is the bending stiffness of the membrane, and *F*_th_ is the tether force^33^. Values of apparent membrane tension were previously reported to be on the order of 0.01 − 1mN/m^19,34,35^. Measurements of tether forces on the unsupported plasma membrane in cellular blebs revealed that membrane tension *γ*_mem_ was contributing a fraction of 30 − 50% percent to measured tether forces, i.e. a fraction of ≈ 10 − 25% of apparent membrane tension^36^.

To determine also changes of apparent membrane tension upon perturbation of cytoskeletal structures, we utilized tether pulling with the atomic force microscope as has been described before, see^22^ and Materials and Methods. In this assay, the AFM cantilever was brought into contact with the cell surface for 1 s. Subsequently, the cantilever is retracted giving first rise to a force decrease down to negative forces that stem from remaining adhesive contact between the cantilever and membrane tethers. Lifting the cantilever further, membrane tethers gradually detach giving rise to sudden force jumps, see red ellipses in Fig. 4a. The magnitude of a measured force jump was interpreted as membrane tether force. In untreated HeLa cells in mitotic arrest, we find typical tether forces of 10 − 30 pN, see Fig. 4b, in agreement with previous tether force measurements^21,24,35,37^. Assuming a typical membrane bending stiffness of *B* = 0.14 pN*µ*m^19,35^, this corresponds to an apparent membrane tension range of ≈ 0.01 − 0.14 mN/m. Measuring tether forces with and without cytoskeletal drugs present, we find that tether forces are significantly reduced upon direct inhibition of myosin with PN-Blebbistatin (Fig. 4c blue violin plot), but not through inhibition of ROCK with Y27632 (Fig. 4c dark red violin plot). Inhibition of actin nucleators formin and Arp2/3 via addition of SMIFH2 and CK666 to the medium leads to a clear reduction of tether forces (Fig. 4c violet and green violin plots) with concomitant induction of cell blebbing reflecting reduced adhesion between plasma membrane and the actin cortex upon treatment, see Supplementary Fig. S3f,g. Inhibition of ERM proteins that can tether the actin cortex to the plasma membrane surprisingly did not trigger a measurable change of tether forces in our assay (Fig. 4c orange violin plot). In accordance with a lack of cell blebbing upon ERM inhibition, see Supplementary Fig. S3h, we conclude from this observation that ERM inhibition does not significantly affect cortex-membrane attachment during mitosis, likely because this function is redundantly supported by other molecules such as integrin complexes and actin-nucleator complexes. Lastly, depolymerization of microtubules through Nocodazole leads to an increase in tether forces (Fig. 4c yellow violin plot). We speculate that the observed effect might emerge indirectly through activation of myosin II upon microtubule disruption^38–41^. We note that, interestingly, the change of apparent membrane tension upon microtubule disruption is opposite to the change of membrane tension as suggested by FliptR fluorescence lifetimes. Finally, we conclude that myosin contractility and actin nucleators contribute to the maintenance of apparent membrane tension.

**Figure 4.**
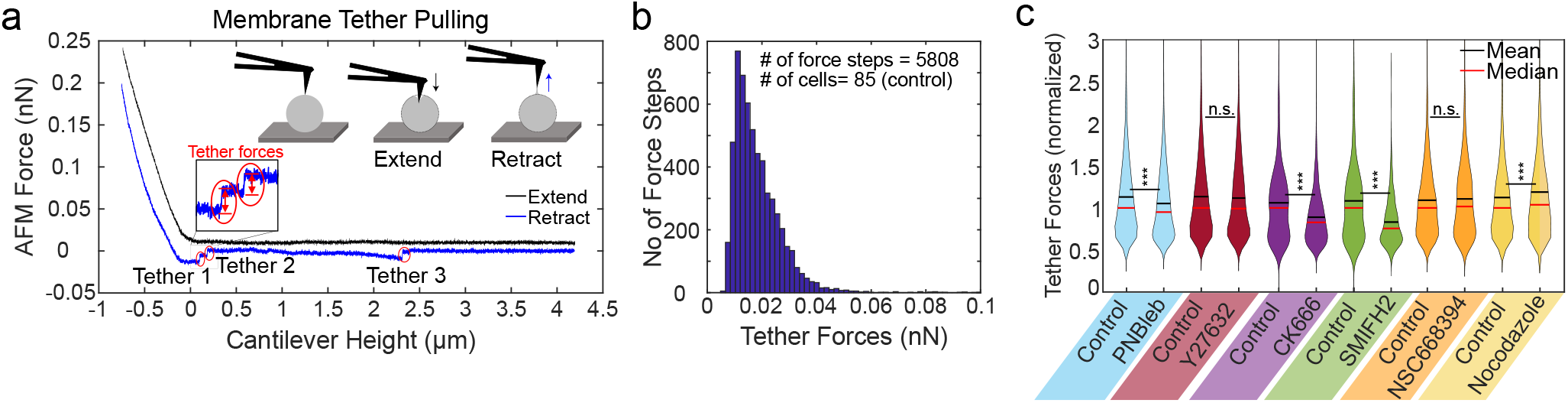
Changes of AFM tether forces upon cytoskeletal drug treatment in mitotic HeLa cells. a) AFM force in dependence of cantilever height during an indentation (black curve) and retraction (blue curve) for an exemplary cell. Force jumps in the retraction force curve are interpreted as tether forces as indicated. b) Histogram showing tether force distribution for untreated cells in mitotic arrest. c) Violin plots showing normalized tether forces for untreated cells and cells treated with cytoskeletal drugs as indicated (concentrations as in Fig. 2). Forces were normalized by the median of the control measurements of the same day. Significance was tested using a two-tailed Mann-Whitney U test and *p* ≤ 0.05. The number of measured cells *n*_*C*_ and the number of measured force steps *n*_*T*_ was blue: control: *n*_*C*_ = 50, *n*_*T*_ = 3953, PN-Blebbistatin: *n*_*C*_ = 50, *n*_*T*_ = 4628, red: control: *n*_*C*_ = 40, *n*_*T*_ = 3540, Y27632: *n*_*C*_ = 40, *n*_*T*_ = 3352, violet: control: *n*_*C*_ = 40, *n*_*T*_ = 1497, CK666: *n*_*C*_ = 40, *n*_*T*_ = 2173, green: control: *n*_*C*_ = 30, *n*_*T*_ = 1087, SMIFH2: *n*_*C*_ = 30, *n*_*T*_ = 958, orange: control: *n*_*C*_ = 40, *n*_*T*_ = 2927, NSC668394: *n*_*C*_ = 40, *n*_*T*_ = 2589, yellow: control: *n*_*C*_ = 35*n*_*T*_ = 2689, Nocodazole: *n*_*C*_ = 35, *n*_*T*_ = 2141.

### E. Impact of cytoskeletal inhibitors on actin cortex tension

So far, our focus has been on measuring membrane tension or apparent membrane tension using FliptR and membrane tether forces. From a biomechanical point of view, also actin cortex tension is a crucial factor influencing membrane tension as it regulates the hydrostatic pressure in the cell that pushes against the plasma membrane at the cell periphery, see Section III and Fig. 7b^11^. To explore the influence of possible cortex tension changes on membrane tension upon application of cytoskeletal inhibitors, we measured the actin cortical tension in HeLa cells in mitotic arrest with and without inhibitors using an AFM-based parallel plate cell confinement assay, see Fig. 5a, Materials and Methods and^30,42,43^. In this setup, a wedged cantilever is utilized to confine mitotic cells between the AFM cantilever and a glass-bottom dish. Cortical tension is calculated from the AFM force required to confine the cell, the confinement height, and the cell diameter during confinement, see Materials and Methods and^42^. In Fig. 5b, we display a boxplot of measured cortical tension values in control conditions (black boxes) and upon cytoskeletal perturbations through addition of indicated drugs to the medium. As expected, we find that the myosin inhibitor PN-Blebbistatin and the ROCK inhibitor Y27632 reduce cortical tension significantly (blue and dark red box). While actin polymerization perturbation by the Arp2/3 inhibitor CK666 did not result in a significant cortical tension change (violet box), inhibition of actin polymerization by formin inhibitor SMIFH2 significantly reduced cortical tension (green box). Further, inhibition of ERM proteins by NSC668394 also resulted in a significant reduction in cortical tension (orange box). In contrast, the microtubule inhibitor Nocodazole led to an increase in cortical tension (yellow box) in accordance with previous findings of myosin II activation upon microtubule disruption^38–41^.

**Figure 5.**
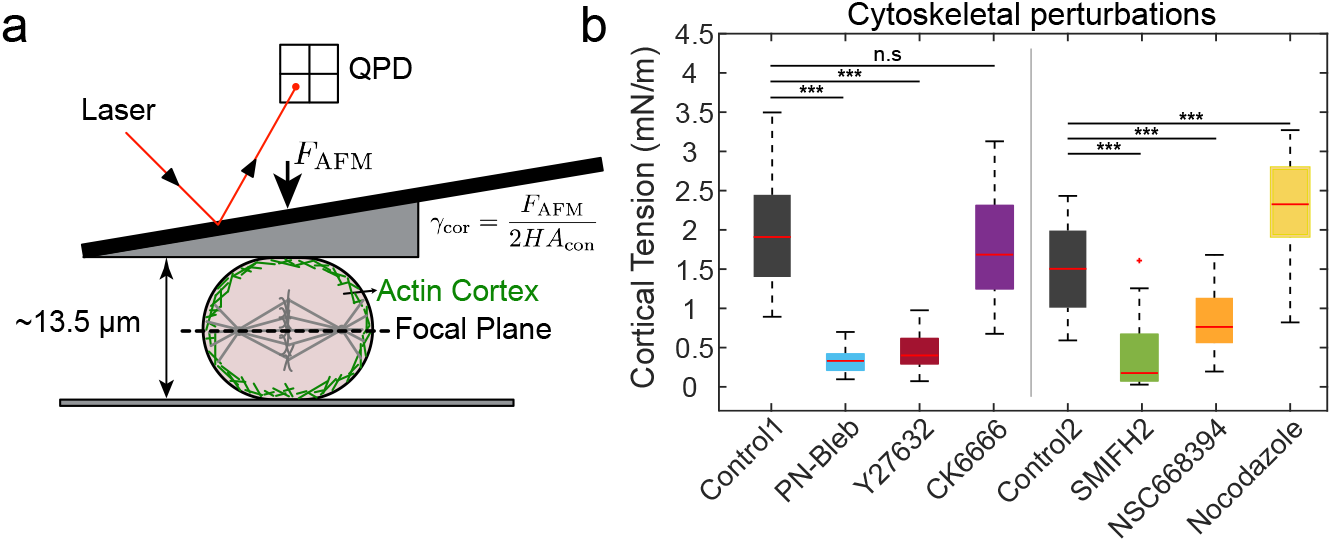
Changes of cortical tension upon cytoskeletal drug treatment. a) Schematics of a mitotic cell confined between a glass coverslip and the wedge of an AFM cantilever. AFM force *F*_AFM_ and the imaging-derived mean curvature *H* and contact area *A* are used to calculate cortical tension *γ*_cortex_, see Materials and Methods. b) Boxplot showing cortical tensions of HeLa cells in mitotic arrest with and without cytoskeletal drugs (concentrations as described in Fig. 2). Cortical tensions in different conditions were compared to control measurements on the same day. Significance was tested using a two-tailed Mann-Whitney U test and *p*≤ 0.05.

### F. Impact of osmotic shock and cytoskeletal perturbations on filopodia

Due to the limited stretchability of plasma membranes (rupture strains of ≈ 3%^1,15^), membrane reservoirs play a major role in the regulation of membrane tension aside from hydrostatic pressure^37^. In mitotic HeLa cells, filopodia constitute a major membrane reservoir as is evident from confocal imaging, see Supplementary movie 1 and Fig. 6b,e,f. To better understand membrane tension changes observed during perturbations applied in our study, i.e. osmotic shock and cytoskeletal perturbations, we decided to quantify the concomitant change in filopodia abundance.

**Figure 6.**
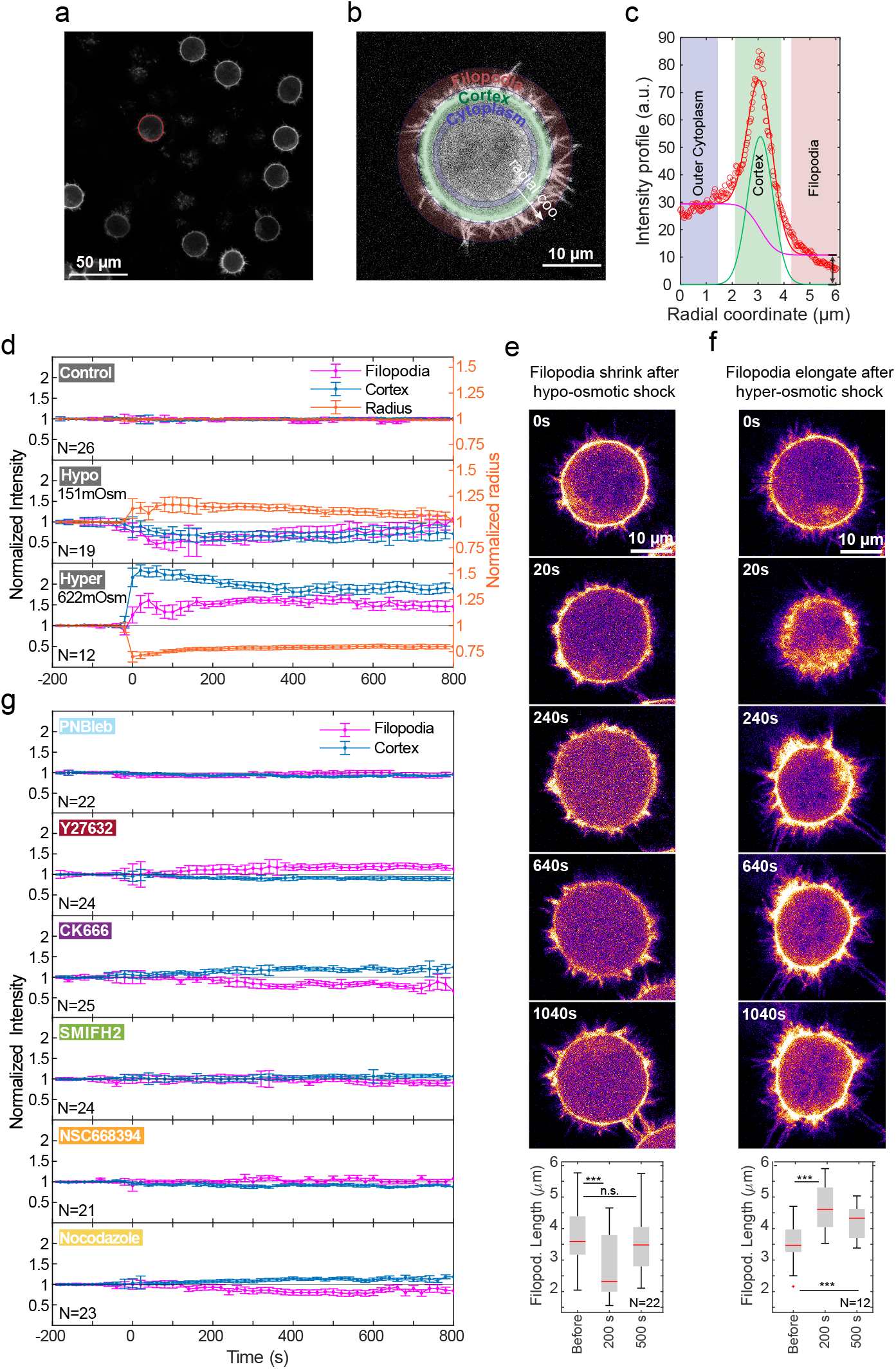
Changes in filopodia abundance in response to osmotic shock and cytoskeletal perturbations via cytoskeletal drugs (concentrations as in Fig. 2). a) Exemplary confocal microscopy image acquired for filopodia quantification in HeLa cells in mitotic arrest. Cells express Lifeact-mScarlet as a label of f-actin. The focal plane was chosen to coincide approximately with the equatorial plane of mitotic cells. The red circle shows the edge detection of the cell boundary of the mitotic cell selected for analysis. (Scale bar: 50 *µ*m.) b) Close-up of one HeLa cell, illustrating the distribution of f-actin fluorescence to filopodia, cortical actin, and cytoplasmic actin based on Lifeact. Correspondingly, a red, green, and blue annulus region highlights the different regions of f-actin intensity. Intensities in each of these regions were quantified over time as described in c. c) Total radial intensity profile of the mitotic cell in panel b (red data points). The solid red line shows a fitted intensity profile obtained by image analysis, see Materials and Methods. The magenta curve reflects the fitted constant cytoplasmic intensity in the outer region of the cytoplasm, which transitions to the fitted constant fluorescence intensity contributed by filopodia outside the cortex. The cortical intensity is captured by the fit of a blue Gaussian curve. The magenta and green curves combined produce the red curve, which closely fits the measured radial intensity profile. d,g) Time evolution of filopodia and cortical intensity during hypo- and hyper-osmotic shock (d) or cytoskeletal perturbations (g) using cytoskeletal drugs (concentrations as in Fig. 2). Perturbations were applied at time point zero. Control measurements were conducted by adding isotonic cell culture medium to assess the effects of triggered fluid fluxes. e,f) Time series of confocal fluorescence microscopy images during osmotic shocks of a single exemplary HeLa cell in mitotic arrest demonstrating the dynamics of filopodia and cell size over time. Boxplots show the decrease (increase) of averaged filopodia length upon hypo(hyper)-osmotic shock. Significance was tested using a two-tailed, paired t-test and *p*≤ 0.05.

**Figure 7.**
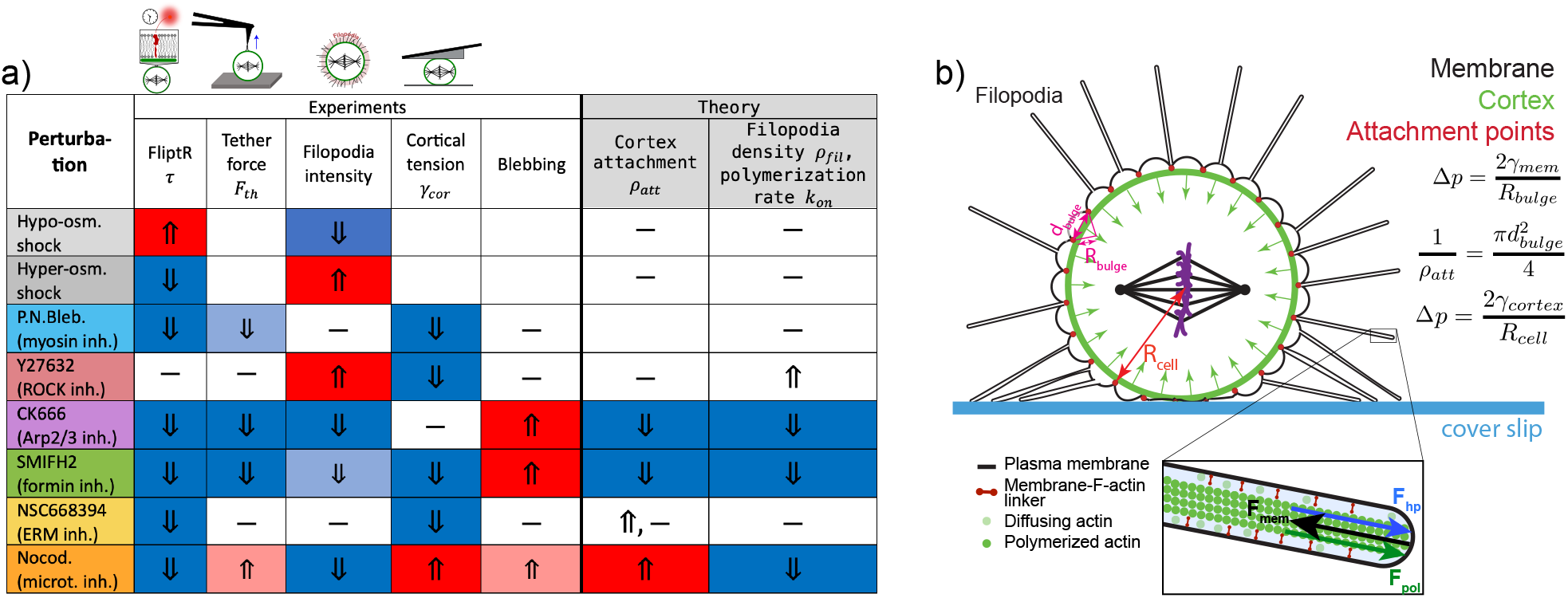
a) Summary of observed experimental changes upon perturbations (blue: decline, red: increase, -: no change, blank: no measurement or no conclusive insight). Right half: Predicted changes of model parameters *ρ*_att_, *ρ*_fil_, and *k*_on_ upon drug treatment. If two symbols are plotted, two possibilities for changes are commensurate with the observations. b) Schematic of membrane structures and force balances inside the mitotic cell. Membrane is either stored in membrane bulges that form over a mesh of membrane-cortex attachment with a certain diameter (*d*_bulge_) due to hydrostatic pressure Δ*p* pushing against the membrane from the inside. Hydrostatic pressure is balanced by both membrane tension (inside a bulge) and by the cortex, such that Laplace’s law is fulfilled with either the curvature radius of the bulge or the cortex, respectively. Bulge diameters are strongly exaggerated in this image for illustrative purposes. The inset shows the force balance between outwards-pushing hydrostatic and actin polymerization pressure and retracting membrane tension at a filopodial tip.

Assuming that the number of actin filaments in filopodia is constant, we infer that the amount of membrane stored in filopodia is proportional to the amount of f-actin in filopodia. To quantify filopodia-associated f-actin as a measure of filopodial membrane reservoir, we used transgenic HeLa cells expressing fluorescently labeled Lifeact - an established label of f-actin^44^. More concretely, we performed confocal imaging of Lifeact in the equatorial plane of HeLa cells in mitotic arrest and quantified Lifeact fluorescence in cellular cross-sections in three radial sectors: the outer cytoplasm, the cortex, and the filopodia region, see Fig. 6a-c and Materials and Methods.

As demonstrated in Fig. 6d,e, both filopodial and cortical f-actin intensities decreased following hypo-osmotic shock applied at time point *t* = 0 s, correlating with an increase in cell radius. Conversely, upon hyper-osmotic shock, cell radii decreased while both cortical and filopodial f-actin intensities increased, see Fig. 6d,f. Further, there is a gradual relaxation of cell radius, filopodia, and cortical f-actin intensity over time, likely due to active cell volume regulation. Concomitant changes in filopodia length, see Fig. 6d,f suggest that the observed changes in filopodia abundance are at least in part attributed to such length changes.

In the context of cytoskeletal perturbations, we observed no change in filopodial f-actin levels upon myosin inhibition with PN-Blebbistatin, see Fig. 6g, 1st panel and Supplementary Fig. S3d. However, ROCK inhibition with Y27632 led to a marked increase in filopodial f-actin (Fig. 6g, 2nd panel and Supplementary Fig. S3e). These findings suggest that filopodial dynamics are not directly influenced by myosin activity but are instead modulated by other downstream effectors of ROCK such as cofilin. Perturbing actin polymerization by the Arp2/3 inhibitor CK666 led to reduced filopodial f-actin. To a lesser extent, this effect was also visible upon perturbation of actin polymerization by the formin inhibitor SMIFH2, see Fig. 6g, 3rd and 4th panel and Supplementary Fig. S3f,g. Performing inhibition of ERM proteins with NSC668394 did not significantly alter filopodial f-actin intensity, see Fig. 6g, 5th panel and Supplementary Fig. S3h. Finally, perturbing microtubule polymerization by Nocodazole treatment, we also observed reduced filopodial f-actin intensity, see Fig. 6g, lowest panel and Supplementary Fig. S3i.

In summary, we compiled the changes in FliptR lifetimes, tether forces, cortical tension, and filopodia abundance upon cytoskeletal perturbations and osmotic shock, as shown in Fig. 7a.

## III. MODEL OF MEMBRANE TENSION GENERATION

We derived a biophysical model of membrane tension regulation to account for our experimental observations. As illustrated in Fig. 7b, we anticipate that the membrane in the mitotic cell is either stored in filopodia or in bulge-like membrane compartments that are directly above the actin cortex. At their periphery, bulges are tethered to the actin cortex. For simplicity, we assume in our theoretical derivations that such bulges have the shape of a spherical cap. Bulges can be more or less inflated depending on their base diameter *d*_bulge_ and the hydrostatic pressure Δ*p* inside the cytoplasm. This hydrostatic pressure Δ*p* is balanced by active cortical tension *γ*_cortex_ at the cell periphery, see Fig. 7b. Denoting the curvature radius of the(bulge wiJth *R*_bulge_ and assum) ing *R*_bulge_ ≥ *d*_bulge_*/*2, we can infer that the bulge area is given by 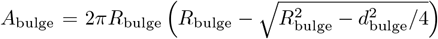 where 1*/*(*π*(*d*_bulge_*/*2)^2^) = *ρ*_att_ with *ρ*_att_ the area density of bulges on the cell surface. Using Laplace’s law, the curvature radius of the bulge *R*_bulge_ can be related to the curvature radius *R*_cell_ of the cortex via *R*_bulge_ = *γ*_mem_*R*_cell_*/γ*_cor_, see Fig. 7b. The total membrane area amounts to

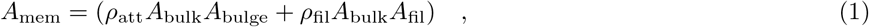

where *ρ*_fil_ is the area density of filopodia and 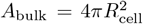 and *A*_fil_ = 2*πR*_fil_*L*_fil_ are the projected surface area of the cell and the average surface area of a filopodium. Here, *L*_fil_ and *R*_fil_ denote the length and the radius of the filopodium, respectively. Note that *ρ*_att_*A*_bulk_ and *ρ*_fil_*A*_bulk_ are the overall number of bulges and filopodia covering the surface of a cell.

### Relation between membrane tension and filopodia length

The concentration profile *c*(*t, x*) of actin monomers along the length of a filopodium has to satisfy the diffusion equation 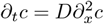, with the boundary conditions *c*(*t*, 0) = *c*_0_ at the base of the filopodium and the diffusive flux − *D∂*_*x*_*c*(*t, L*_fil_) = *k*_*on*_*c*(*t, L*_fil_) at the tip of the filopodium where *k*_on_*c*(*t, L*_fil_) is the polymerization rate at the tip and *x* is the position coordinate along the length of the filopodium. Here, *c*_0_ is the actin monomer concentration in the cytoplasm and *D* is the corresponding diffusion coefficient. In stationary state, we have that *∂*_*t*_*c* = 0 and thus we find that 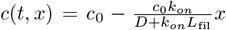. We assume now that the polymerization force of actin filaments at the tip of the filopodium is given by *F*_*pol*_ = *ξk*_*on*_*c*(*t, L*_fil_), where *ξ* is a proportionality constant. Assuming a force balance *F*_*pol*_ + *F*_*hp*_ = *F*_mem_, where 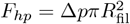 is the outwards-directed force due to hydrostatic pressure and *F*_mem_ = 2*πR*_fil_*γ*_mem_ the retracting force from membrane tension (Fig. 7b), we obtain

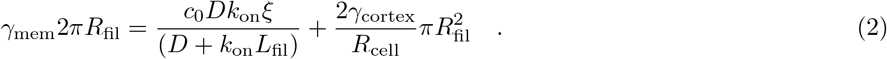

We anticipate a bulk concentration *c*_0_ of actin monomers available for polymerization as *c*_0_ ≈ 2.5 × 10^3^ *µ*m^−3^, i.e. about 5% of the previously estimated concentration of total actin, see^45^. Estimating the retrograde flow within filopodia as *v*_retr_ ≈ 25 nm/s, see^46^, the actin monomer diameter as *d*_mon_ ≈ 5 nm see^28^, the number of actin polymers within a filopodium as *N*_pol_ ≈ 20, see^47^, and the radius *R*_fil_ as 0.1 *µ*m, see^47^, we can estimate *k*_on_*c*(*L*_fil_) through the mass conservation relation 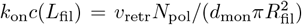. Further, upon mechanical rupture of their tip connection, filopodia exert a passive retraction force of *F*_retr_ = 15 pN, see^46^. With force balance *F*_retr_ = *ξk*_*on*_*c*(*L*_fil_) at the tip and the above estimates for *k*_on_*c*(*L*_fil_), we obtain *ξ* ≈ 5 × 10^−6^ nN *µ*m^2^s.

### Membrane tension is regulated through cortical tension and actin polymerization rates

Assuming that total membrane area *A*_mem_ is constant, we can calculate *γ*_mem_ and *L*_fil_ from Eqn. (1) and (2), provided that cell radius *R*_cell_, cortical tension *γ*_cortex_, cortex-membrane attachment density *ρ*_att_ and filopodial density *ρ*_fil_ are known, see Fig. 8 and Materials and Methods. Assuming *ρ*_att_ = 10 − 100 *µ*m^−2^, see^48^, *γ*_cortex_ = 0 − 2.5 mN/m, see Fig. 5, *ρ*_fil_ = 1 *µ*m^−2^ and 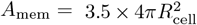 (rough estimate from filopodia imaging, see Supplementary Movie 1), we find membrane tensions in the range 0.02 0. − 08 mN/m for a typical cell radius *R*_cell_ = 10 *µ*m. These tension values are commensurate with previous measurements^33^. Corresponding model predictions of filopodia lengths range between ≈ 2 − 6 *µ*m and are therefore comparable to observed filopodia lengths, see Fig. 8b and 6b. As expected, cortical tension increases membrane tension, see Fig. 8a, by increasing hydrostatic pressure. At the same time, the height of the membrane bulges increases, see Fig. S4. By contrast, cortex-membrane attachment barely affects membrane tension, see Fig. 8a. Increased density of filopodia increases membrane tension, due to reduced emergent filopodia lengths and correspondingly increased filopodial actin polymerization forces because of higher actin monomer concentrations at the filopodia tips, see Fig. 8a,b. By the same mechanism, increase of *k*_on_ (or alternatively bulk actin monomer concentration *c*_0_) leads also to an increase of membrane tension and increased filopodia length, see Fig. 8c,d.

**Figure 8.**
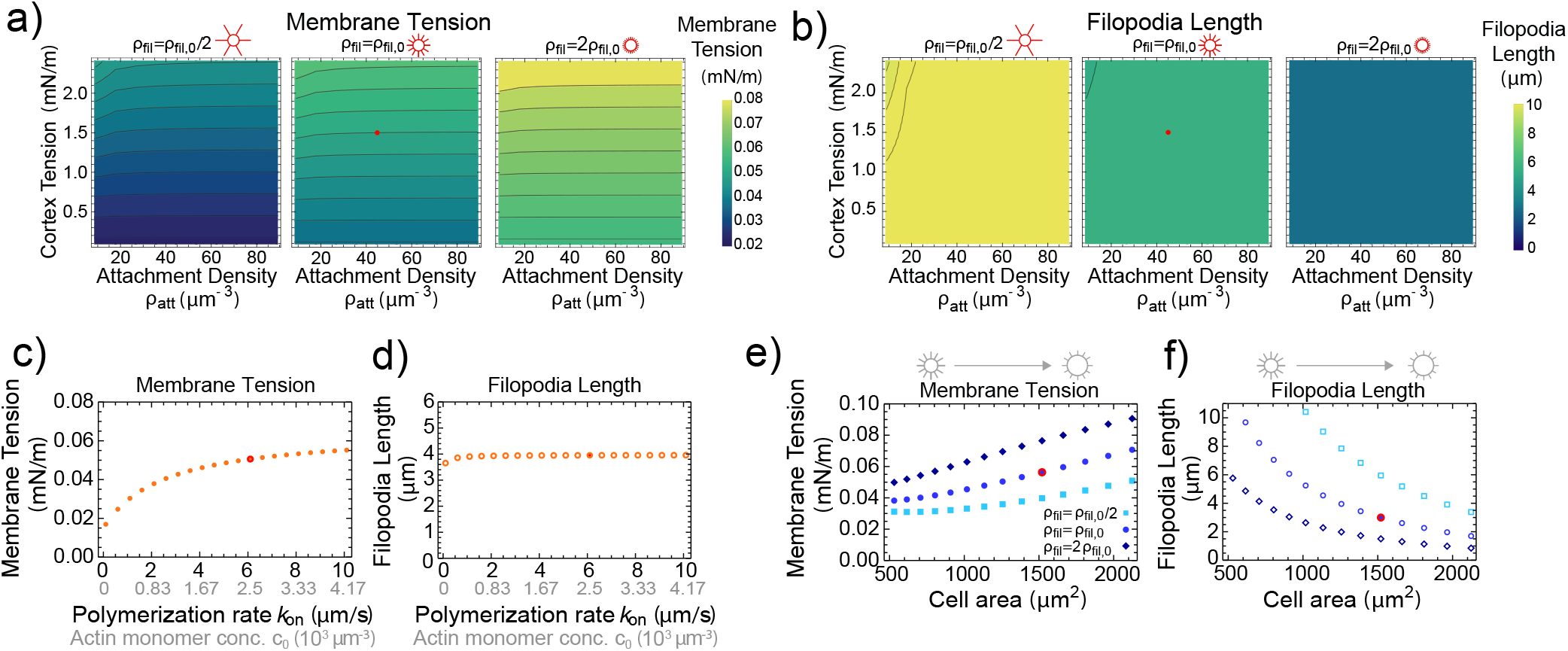
Model prediction of membrane tension and filopodia length. a,b) Membrane tension and filopodia length as a function of cortical tension *γ*_cortex_, cortex-membrane attachment density *ρ*_att_ and area density of filopodia *ρ*_fil_. From left to right, filopodial density increases (Left: *ρ*_fil_ = *ρ*_fil,0_*/*2, Middle: *ρ*_fil_ = *ρ*_fil,0_, Right: *ρ*_fil_ = 2*ρ*_fil,0_ with *ρ*_fil,0_ = 0.25 *µ*m^−2^). c,d) Membrane tension and filopodia length increase with filopodial actin polymerization rate *k*_on_ (with actin monomer concentration *c*_0_). e,f) Membrane tension and filopodia length as a function of varying cell area upon cell swelling or shrinkage. If not indicated otherwise, parameters are set to our choice of reference parameters: concentration of free actin monomers *c*_0_ = 2.5 *×* 10^3^*µ*m^−3^, *k*_on_ = 6 *µ*m*/s, ξ* = 5*×*10^−6^ nN · *s* · *µ*m^2^, *D* = 10 *µ*m^2^*/s, R*_fil_ = 0.1 *µ*m, *ρ*_fil_ = 1 *µ*m^−2^, *ρ*_att_ = 44.44 *µ*m^−2^, *R*_cell_ = 10 *µ*m and total membrane area 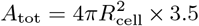. In all panels, data points that correspond to reference parameters are indicated with red marks.

### Model predictions upon osmotic shock

Finally, to better understand the effect of hypo-osmotic shock within the scheme of the model, we calculated changes of membrane tension upon an increase of the cell radius. Increased storage of membrane at the cell periphery due to the increased projected surface area of the cell leads to decreased filopodia lengths. Due to higher polymerization power of shorter filopodia, membrane tension is increased jointly, see Fig. 8e,f. Correspondingly, model predictions agree with our experimental observations, compare Fig. 1g and Fig. 6d,e. Conversely, decreasing cell radii as during a hyper-osmotic shock, filopodia increase in length and membrane tension declines, see Fig. 8e,f. Again, these model predictions are in accordance with experiments, compare Fig. 1g and 6d,f.

## IV. DISCUSSION

In this study, we examined the influence of the cytoskeleton on plasma membrane tension. HeLa cells arrested in mitosis were selected as the biological model system due to their well-defined, rounded shape, which facilitates the analysis of the plasma membrane independently from internal membrane structures and cell cycle-induced changes. Our primary focus was on the role of the actin cortex in regulating plasma membrane tension, given its direct interaction with the membrane. Additionally, we investigated the contribution of filopodia and microtubules.

Plasma membrane tension changes induced by cytoskeletal perturbations were measured using the membrane-associated dye FliptR, which reports variations in membrane tension through shifts in its fluorescence lifetime^19,23–27^. The reliability of this probe in our experimental setup was confirmed through osmotic shock experiments (see Fig. 1). To investigate the specific contributions of cytoskeletal components to membrane tension, we introduced acute perturbations using targeted cytoskeletal drugs. Inhibiting myosin, Arp2/3, formins, ERM proteins, and microtubules through pharmacological inhibitors (PN-Blebbistatin, CK666, SMIFH2, NSC668394, and Nocodazole), we find an emerging reduction of FliptR lifetimes indicating a decline of membrane tension. Interestingly, inhibition of the ROCK inhibitor Y27632 did not change FliptR lifetimes. Furthermore, we assessed changes in apparent membrane tension by measuring membrane tether forces under conditions with and without cytoskeletal drugs (see Fig. 2). In this assay, we found that pharmacological inhibition of myosin, Arp2/3 and formins reduced tether forces. By contrast, microtubule disruption elevated tether forces, while ROCK and ERM inhibition did not induce any significant changes.

To gain a better mechanistic understanding of how the cytoskeleton affects membrane tension, we also measured concomitant changes of critical biophysical factors such as actin cortex tension, see Fig. 4. As expected, actin cortex tension reduced upon myosin and ROCK inhibition with PN-Blebbistatin and Y27632, respectively. Furthermore, inhibition of formins and ERM proteins via inhibitors SMIFH2 and NSC668394, respectively, markedly reduced cortical tension - both in agreement with earlier reports on mitotic cortex tension regulation^43,48,49^. Inhibiting Arp2/3-mediated actin nucleation did not significantly affect cortical tension, consistent with the unchanged cortex thickness observed in mitotic HeLa cells upon Arp2/3 inhibition^48^. Disruption of microtubules with Nocodazole was the only treatment that led to increased cortical tension as has been reported before^32^.

As a second critical biophysical factor for membrane tension regulation, we investigated membrane reservoirs stored in filopodia. As has been reported before^50–52^, filopodial membrane reservoirs are very abundant in mitotic cells storing excess membrane released during mitotic rounding, see also Fig. 6 and Supplementary movie 1. Upon application of cytoskeletal drugs, we partly observed changes in filopodial abundance judged by f-actin fluorescence intensity, see Fig. 6. In particular, treatment with the ROCK inhibitor Y27632 enhanced filopodia formation, while actin polymerization inhibitors (SMIFH2 and CK666) reduced filopodia levels. Moreover, microtubule disruption upon Nocodazole treatment reduced filopodia formation, suggesting an indirect effect of microtubules on actin polymerization dynamics. By contrast, inhibition of myosin and ERM proteins did not induce any changes.

To account for our observations, we developed a mathematical model that describes how the plasma membrane tension attached to a contractile actin cortex is regulated in the presence of numerous filopodial structures. At the core of this model is a force balance between hydrostatic pressure and contractile cortical tension at the cell periphery, as well as between hydrostatic pressure and plasma membrane tension, see Fig. 7b. Furthermore, we assumed a force balance at the tip of filopodia that incorporates a retracting force from membrane tension and a protruding force from hydrostatic pressure and actin polymerization, see Eq. (2). The model predicts that in such a system, membrane tension is driven by two main factors: i) pressure-generating cortical tension and ii) actin polymerization against the plasma membrane inside filopodia. Correspondingly, an isolated change in cortical tension would trigger a corresponding change in membrane tension. On the other hand, an isolated change in filopodial abundance or filopodial actin polymerization rate would trigger a corresponding change in membrane tension without changes of hydrostatic pressure. Membrane storage in sparse but long (rather than short and abundant) filopodia tends to reduce membrane tension due to depleted actin monomer concentration and polymerization rates at filopodial tips. Further, our experiments put forward that both membrane tension generation through filopodia and hydrostatic pressure rely on free unsupported cell surfaces that are not in contact with a substrate and can thus build up curvature. Therefore, contact area increase leads to reduced membrane tension, see Fig. 3. For apparent membrane tension measured through tether forces, changes in cortex-membrane attachment density are predicted to be a critical factor, while the ‘bare’ membrane tension remains largely unaffected by this parameter (see Fig. 8).

Leveraging insights from our model and observed alterations in cortical tension and filopodia dynamics, we can interpret observed FliptR lifetime and membrane tether force changes upon cytoskeletal perturbations, see Fig. 1 and Fig. 2; upon myosin inhibition, we observed reduced FliptR lifetimes and tether forces which we attribute to reduced membrane tension due to lowered cortical tension and hydrostatic pressure inside the cell. By contrast, ROCK inhibition did neither change membrane tension nor apparent membrane tension. While this treatment also reduces cortical tension and hydrostatic pressure just like myosin inhibition, we also found increased filopodia abundance which likely acts antagonistically on membrane tension leaving membrane tension overall unchanged. Inhibition of actin nucleators Arp2/3 and formin led to a decrease in FliptR lifetimes and tether forces suggesting an induced decline of membrane tension and apparent membrane tension (Fig. 2). We attribute the decline in membrane tension to filopodia reduction upon both treatments and, in the case of formin inhibition, in addition to declined cortical tension. Both treatments were accompanied by increased cell blebbing which lead us to conclude that declined tether forces are promoted by reduced cortex-membrane adhesion. Upon pharmacological inhibition of ERM proteins, we found reduced FliptR lifetimes but unchanged tether forces suggesting reduced membrane tension but unchanged apparent membrane tension. While the former can be accounted for by a concomitant decline in cortical tension and hydrostatic pressure upon ERM inhibition, see Fig. 5, unchanged tether forces suggest that membrane-cortex attachment is unchanged or even slightly increased upon NSC668394 treatment in mitosis. In conjunction with a lack of cell blebbing upon treatment, this suggests that pERM proteins are dispensable for cortex-membrane attachment perhaps due to their preference for positive membrane curvature^53^. Finally, we find that also microtubules seem to play a regulatory role in mitotic membrane tension. Upon microtubule disruption, FliptR lifetimes are reduced suggesting a reduction in membrane tension *γ*_mem_ likely due to a decrease in filopodia abundance in spite of a slight cortical tension increase. Interestingly, tether forces are increased hinting at a rise in apparent membrane tension *γ*_app_ = *γ*_adh_ + *γ*_mem_ upon microtubule disruption. We conjecture that an over-proportional increase of cortex-membrane attachment *γ*_adh_ possibly related to RHOA-ROCK activation^40^ is at the heart of this effect.

Based on our findings, we put forward that plasma membrane tension in cells is regulated by two primary factors: hydrostatic pressure inside the cytoplasm generated by in-plane cortical tension at the cell periphery as well as actin polymerization forces orthogonal to the cell periphery e.g. inside filopodial tips. In particular, our study highlights the fact that it is not the amount of membrane stored in membrane reservoirs as such that regulates membrane tension but the strength and density of forces transmitted to the membrane. Further, we find that the apparent membrane tension, which is measured using tether forces, is not a reliable indicator of the evolution of ‘bare’ membrane tension due to its dependence on membrane-cortex adhesion.

## V. MATERIALS AND METHODS

### A. Cell Culture

In this study, two types of cells were utilized: HeLa Kyoto and a transgenic HeLa Kyoto cell line that expresses Lifeact-mScarlet (Addgene #85054), see^30^. The pLifeAct-mScarlet-N1 vector was obtained as a gift from Dorus Gadella (Addgene plasmid #85054; http://n2t.net/addgene:85054; RRID: Addgene 85054). HeLa Kyoto cells were cultured in Dulbecco’s Modified Eagle Medium (DMEM) supplemented with 10% fetal bovine serum (FBS) and 1% penicillin/streptomycin. For the transgenic HeLa cells, 0.4 mg/mL of geneticin was additionally included in the culture medium. Cell culture flasks were maintained at 37°C in a 5% CO_2_ environment. Cells were passaged every 2 − 3 day when they reached 60-80% confluency.

### B. FliptR lifetime measurements in combination with osmotic shock or drug treatments

One day before the experiments, 20, 000 cells were seeded into an Ibidi *µ*M-Slide 8-Well glass bottom dish (1 cm^2^ growth area, Cat. No. 80827). As previously described, 2 to 8 hours before measurement, the growth medium was replaced with a CO_2_-independent imaging medium, and STC (2 *µ*M) (#164739, Sigma) was added to arrest the cells in mitosis. FliptR dye was introduced at a concentration of 1 *µ*M at least 15 minutes before the start of the measurement.

The experiments were conducted using an LSM 780 confocal microscope (Zeiss) at the CMCB light microscopy facility at TU Dresden. A Plan Apochromat 20 × */*0.8 air objective (Zeiss) and a 20 MHz pulsed 473 nm excitation laser at 33% power were utilized. During the measurements, the cells were maintained at 37°C inside an incubation chamber. For each measurement, a region of interest with more than 10 mitotic cells was chosen. The focus was adjusted to the equatorial cross-section of the mitotic cells. The imaging settings were as follows: bi-directional scanning, 512 × 512 pixels^2^, pixel size of 0.83 *µ*m, frame averaging of 8 to 16 frames, pinhole size of 2.46 Airy units, pixel dwell time of 1.27 *µ*s, and a time series with a 20 s interval. FLIM images were captured as “.sdt” images in FIFO Imaging mode. In total, 60 to 70 cycles were recorded with a collection time of 10 s (repetition time: 20 s, display time: 20 s, gain: 2).

After approximately 10 cycles of scanning, either osmotic shock or cytoskeletal drug addition was performed by adding medium or water with and without cytoskeletal drugs to the final concentrations as indicated. Afterwards, the effects of the perturbation were observed over the next 50 to 60 cycles. For a 50% v/v hypo-osmotic shock (199 mOsm), 100 *µ*L of distilled water was added to 200 *µ*L of cell medium, while for a 100% v/v hypo-osmotic shock (151 mOsm), 200 *µ*L of distilled water was added to 200 *µ*L of cell medium. To induce hyperosmotic shock, sorbitol solutions at concentrations of 1000 mM and 2000 mM were prepared in the imaging medium. During the experiment, hyperosmotic conditions of 50 mM (348 mOsm), 75 mM (365 mOsm), 622 mOsm, and 938 mOsm were achieved by adding 10.53 *µ*L, 16.2 *µ*L, 100 *µ*L of a 1000 mM sorbitol solution, and 100 *µ*L of a 2000 mM sorbitol solution to 200 *µ*L of the medium, respectively. We pipetted three times to mix the newly added supplemented medium with the already present medium in the well. The osmolarities of the final solutions were determined using a freezing point depression osmometer (Vogel OM818).

To monitor changes following the addition of cytoskeletal drugs, drug solutions were prepared at three times the final concentration in the medium beforehand. During the experiment, 100 *µ*L of this higher concentration solution was added to 200 *µ*L of growth medium in the well to achieve the final concentrations: Y27632 (Rock Inhibitor, 25 *µ*M, #1005583, Cayman Chemical), PN-Blebbistatin (10 *µ*M, #DR-N-111 Optopharm), SMIFH2 (40 *µ*M, #S4826-5MG Sigma), CK666 (50 *µ*M, #29038, Cayman Chemical), Nocodazole (15 *µ*M, #31430-18-9 Sigma), and NSC668394 (50 *µ*M, #BYT-ORB1699797 Biozol).

Data analysis was carried out using a custom-made Matlab code. Respective Matlab codes can be found at https://gitlab.com/polffgroup/FlimAnalysisCortex. In summary, the cell periphery of individual cells was identified as an annular region of high FliptR fluorescence intensity with a width of ≈ 40% of the cell radius centered at the cell periphery. By focusing solely on the lifetimes of photons from this cortical region, FLIM lifetime data were analyzed for individual cells over time. Consequently, the time dynamics of fitted FliptR lifetimes and cell radii following osmotic shock or pharmacological treatment were monitored over time.

### C. AFM in conjunction with FliptR lifetime measurements experiment

One day before the experiments, 10, 000 cells were seeded into a silicon cultivation chamber (growth area: 0.56 cm^2^, ibidi 12-well chamber) attached to 35 mm glass-bottom dishes (FD35-100, Fluorodish). The silicon inserts were removed 2 − 8 hours before measurements, and the growth medium was replaced with a CO_2_-independent imaging medium composed of DMEM (12800-017, Invitrogen), supplemented with 4 mM NaHCO3, buffered with 20 mM HEPES/NaOH (pH7.2) and 10% FBS. At this stage, S-trityl-L-cysteine (STC) (#164739, Sigma) was added to the medium at a concentration of 2 *µ*M to arrest the cells in mitosis. The FliptR dye (1 *µ*M, Tebubio #SC020) and, if applicable, the ROCK inhibitor Y27632 (10 mM, #1005583, Cayman Chemical) were added to the cells at least 15 minutes before the measurement began. Measurements were performed using an inverted LSM780 confocal microscope (Zeiss) from the CMCB light microscopy facility at TU Dresden in combination with an AFM Nanowizard 1 (JPK Instruments). The experiments utilized a Plan Apochromat 20 × */*0.8 air objective (Zeiss) and a 20 MHz pulsed 473 nm excitation laser. The gain in the Becker & Hickl FLIM software was set to 2, providing a time resolution of 0.093 ns for the lifetime detection of collected photons, with a total detection time window of approximately 25 ns. A GFP-RFP filter was used for FLIM imaging. FLIM images of a cell were recorded as ‘.sdt’ files in FIFO Imaging mode, where one file was generated for each time point in the time series. The total cycle number was set to 60 − 80, with a collection time of 4 s, repetition time of 4 s, and display time of 4 s. The confocal microscope acquisition (Zeiss software) was configured for bi-directional continuous scanning of the equatorial plane of a mitotic cell, using frames of 128×128 pixel^2^, with a pixel size of 0.415 *µ*m, frame averaging of 4, a pinhole size of 1.63 Airy units, and a pixel dwell time of 2.54 *µ*s. During the measurements, cells were maintained at 37°C using a Petri dish heater (JPK Instruments). Tipless AFM cantilevers (HQ: CSC37/tipless/No Al, nominal spring constant of 0.3 − 0.8 N/m, Mikromasch) were prepared for the parallel confinement assay by adding a wedge made of UV-curing adhesive (Norland Optical Adhesive 63, Norland Products) to correct their 10° tilt. On each measurement day, the spring constant of the cantilever was calibrated using thermal noise analysis (JPK-SPM software).

Cells were measured sequentially. First, a mitotic cell in the dish was selected based on its shape. An approach was performed with the cantilever on the dish bottom near the chosen cell to determine the relative z-position of the dish bottom. The cantilever was then retracted by about 14.5 *µ*m and gently brought over the mitotic cell to achieve its parallel plate confinement. The force exerted by the cell on the cantilever was allowed to relax before the compression step.

Subsequently, confocal FLIM imaging of the equatorial plane of the selected cell was set up by focusing on the equatorial plane and initiating the FLIM measurement time series in the Becker & Hickl software as described above. The process of uniaxial cell compression was then initiated in the AFM software by lowering the wedged cantilever by 1.5 *µ*m at a speed of 0.2 *µ*m*/*s. The cantilever was held in this lowered position for 100 s, then lifted back to the original height and maintained for another ≈ 50 s before concluding the measurement.

During the measurements, the force on the cantilever and its height relative to the dish bottom were continuously recorded using software from the AFM manufacturer (JPK Instruments). These data, along with the FLIM and confocal images, were used to determine the cell’s surface area, volume, and cortical tension as previously described^30^. The plasma membrane of the cell was identified as an annular region of high peripheral fluorescence intensity. The lifetime of photons from this region was determined by a mono-exponential tail fit of the lifetime histogram, fitting an ≈ 20 ns interval that started 2.5 ns behind the peak of the lifetime histogram. The respective Matlab codes can be found at https://gitlab.com/polffgroup/flimfitting_AFM_FliptR_in_cells.

### D. Cortical tension measurement using AFM

One day before the experiments, 10,000 cells were seeded into a silicon cultivation chamber (growth area: 0.56 cm^2^, from ibidi 12-well chamber) attached to 35 mm glass-bottom dishes (FD35-100, Fluorodish). The silicon inserts were removed 2 − 8 hours before the measurements, and the growth medium was replaced with a CO_2_-independent imaging medium composed of DMEM (12800-017, Invitrogen) with 4 mM NaHCO3, buffered with 20 mM HEPES/NaOH at pH 7.2, and supplemented with 10% FBS. At this stage, S-trityl-L-cysteine (STC) (#164739, Sigma) was added to the medium at a concentration of 2 *µ*M to arrest the cells in mitosis. The cytoskeletal drugs were introduced to the cells at least 15 minutes before the measurement began. Measurements were performed using an inverted LSM510 confocal microscope in combination with an AFM Nanowizard 1 (JPK Instruments). The experiments utilized a Plan Apochromat 20 × */*0.8 air objective (Zeiss). During the measurements, the cells were maintained at 37°C using a Petri dish heater (JPK Instruments). Tipless AFM cantilevers (HQ: CSC37/tipless/No Al, nominal spring constant of 0.3 − 0.8 N/m, Mikromasch) were prepared for the parallel confinement assay by adding a wedge made of UV-curing adhesive (Norland Optical Adhesive 63, Norland Products) to correct for a 10° tilt. On each measurement day, the spring constant of the cantilever was calibrated using thermal noise analysis (JPK-SPM software).

Cells were measured sequentially. First, a mitotic cell was selected based on its shape. Then, an approach was made with the cantilever to the bottom of the dish, in proximity to the chosen cell, to determine the relative z-position of the dish bottom. The cantilever was subsequently retracted by approximately 14.5 *µ*m and gently positioned over the mitotic cell to achieve parallel plate confinement. The force exerted by the cell on the cantilever was allowed to relax before commencing the compression step. The step for uniaxial cell compression was initiated in the AFM software, lowering the wedged cantilever by 1 *µ*m at a speed of 0.2 *µ*m*/*s. The cantilever was maintained in the lowered position for 100 s before being raised back to its original height, allowing time for the AFM force to equilibrate before stopping the measurement.

During the measurements, the force on the cantilever and its height relative to the dish bottom were continuously recorded using software provided by the AFM manufacturer (JPK Instruments). These data were then used alongside the confocal images to determine the cell’s surface area, volume, and cortical tension as previously described^30^. Respective Matlab codes can be found on https://gitlab.com/polffgroup/flimfitting_AFM_FliptR_in_cells.

### E. Filopodia Imaging

To image filopodia, 20, 000 transgenic HeLa Kyoto cells expressing Lifeact-mScarlet were seeded into an Ibidi *µ*-slide 8-well glass bottom dish (1 cm^2^ growth area, Cat. No. 80827) one day before the measurement. As previously described, 2 to 8 hours before imaging, the growth medium was replaced with a CO_2_-independent imaging medium containing 2 *µ*M STC (#164739, Sigma) to arrest the cells in mitosis.

Confocal images of the equatorial plane of mitotic cells were acquired using a Plan-Apochromat 20 × */*0.8 air objective on a Zeiss confocal microscope. The imaging was performed with a continuous 561 nm laser for 60 cycles at 20 s intervals. The imaging parameters were set as follows: unidirectional scanning, frame averaging of 2, an image size of 2048 × 2048 pixels^2^ with a pixel size of 0.1 *µ*m, a pixel dwell time of 0.39 *µ*s, and a pinhole size of 1 Airy unit.

#### Addition of cytoskeletal drugs

To observe changes in the amount of filopodia over time in the equatorial plane, various cytoskeletal drugs were added around the 10th imaging cycle to monitor subsequent changes in f-actin within the cortex and filopodia. To prepare for the experiment, drug solutions were made at three times the required final concentration. During the imaging, 100 *µ*L of this concentrated drug solution was added to 200 *µ*L of growth medium in the well to achieve the specified final concentration; see also Section V B.

#### Filopodia and cortex analysis

Cortex intensity profiles were analyzed as described before, see^31,43^. In short, the cell boundary was identified via intensity thresholding and fitting of a circle using a MATLAB custom code. Along the fitted cell boundary, 200 radial, equidistant lines were determined extending 3 *µ*m to the cell interior and 3 *µ*m into the exterior. The radial fluorescence profiles corresponding to these lines were averaged over all 200 lines. This averaged intensity profile (see Fig. 6c) is then fitted by a linear combination of an error function with negative amplitude (cytoplasmic contribution) and a Gaussian curve (cortical contribution). The respective fit formula is given by

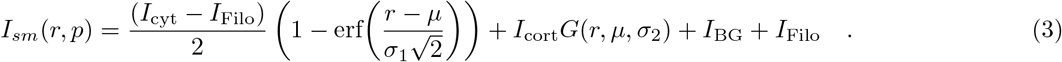

Here *p* = {*µ, σ*_1_, *σ*_2_, *I*_cyt_, *I*_cort_, *I*_Filo_} are fit parameters, which determine the position of the cortex (*µ*), the slope of the error function decay (*σ*_1_), the width of the cortical Gaussian peak *σ*_2_, the amplitudes of the error function and the Gaussian peak (*I*_cyt_ and *I*_cort_) and the intensity contribution of filopodia (*I*_Filo_). *I*_BG_ is the background intensity which is determined as the median intensity value of the entire recorded frame.

By construction, the intensity value converges to (*I*_BG_ + *I*_Filo_) in the cell exterior region and to (*I*_cyt_ + *I*_BG_) in the cell interior. To calculate the cortex intensity, the fitted Gaussian is integrated to give 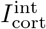 ^31^. Filopodia intensity is quantified by *I*_Filo_.

In Fig. 6d,e, *I*_cort,int_ and *I*_Filo_ are plotted normalized by their initial value (median of the first 8 time points). Error bars in Fig. 6d,e show standard errors of the median, i.e. 1.25 times the standard error of the mean. Respective Matlab codes can be found at https://gitlab.com/polffgroup/filopodiaanalysis.

#### Filopodia Imaging (z-stack imaging)

The 3D structure of f-actin in individual mitotic cells was captured using a Zeiss confocal microscope equipped with a Plan-Apochromat 40 × */*1.0 DIC M27 water objective. We used a 561 nm laser for excitation during the imaging process. Z-stacks of Lifeact-mScarlet fluorescence in the mitotic cells were acquired at a resolution of 2048 × 2048 pixel^2^, utilizing a bottom-to-top z-stack with0.25 *µ*m intervals. The scan area was configured to 42.5 × 42.5 *µ*m^2^.

Using the 3D projection feature of ImageJ (Image/Stacks/3D Projection) with brightest point projection, we combined the acquired z-stack to create a movie of a HeLa cell expressing Lifeact-mScarlet, rotating around the y-axis. The full 360^*°*^ rotation was divided into 36 equidistant segments.

#### Filopodia imaging after contact establishment/removal

Three times series of z-stacks where recorded in conjunction with a dynamic AFM confinement (z-interval: 0.63 *µ*m, time interval: 10 s, pixel size: 0.09 nm) with a Plan-Apochromat 20 × */*0.8 air objective on a Zeiss LSM900 confocal microscope in combination with a Bruker Nanowizard 4 XP at the PoL microscopy facility, TU Dresden. Z-stacks were centered around the top contact plane. The imaging was performed with a continuous 561 nm laser to record fluorescence of Lifeact-mScarlet in selected mitotic HeLa cells. The first time series was recorded at shallow confinement, see Fig. S2a, for a duration of 5 frames. Since there was no visible dynamics in filopodia, we used this initial imaging sequence to estimate the bleaching rate during imaging, see see Fig. S2c. The second time series was started directly after the cantilever had been lowered to reach increased contact area to capture filopodia dynamics within at least 150 s after establishment of contact increase, see Fig. S2a,d. Prior to imaging start, the focal plane was manually lowered by 3 *µ*m to move the center of the focal plane down in conjunction with the top contact area. The third time series was started directly after the cantilever had been retracted thereby reducing contact area again. The goal was to capture filopodia dynamics within at least 50 s after establishment of contact increase, see Fig. S2a,e. Prior to imaging start, the focal plane was manually raised by 3 *µ*m to move the center of the focal plane upwards in conjunction with the top contact area.

### F. AFM membrane Tether pulling

For the AFM membrane tether-pulling experiment, 20, 000 cells were seeded into a silicon cultivation chamber (growth area: 0.56 cm^2^, from the ibidi 12-well chamber) attached to 35 mm glass-bottom dishes (FD35-100, Fluorodish). The silicon inserts were removed 2 to 8 hours before the measurement, and the growth medium was replaced with a CO_2_-independent imaging medium composed of DMEM (Invitrogen) with 4 mM NaHCO3, buffered with 20 mM HEPES/NaOH at pH 7.2, and supplemented with 10% FBS. At this stage, 2 *µ*M S-trityl-L-cysteine (STC) (#164739, Sigma) was added to the medium to arrest the cells in mitosis. For drug-treated conditions, drugs were added at specified concentrations to the fluorodishes and the dishes were incubated in a CO_2_-independent environment at 37°C for 15 to 20 minutes before measurements began.

Measurements were performed with an AFM Nanowizard 4XP (Bruker) mounted on an inverted LSM700 confocal microscope (Zeiss) at the PoL microscopy facility, TU Dresden. The experiments used a Plan Apochromat 20 × */*0.8 air objective (Zeiss) to locate the cells with transmitted light imaging.

Measurements were conducted using an AFM Nanowizard 4XP (Bruker) mounted on an inverted confocal microscope (Zeiss) located at the PoL microscopy facility, TU Dresden. A Plan Apochromat 20 × */*0.8 air objective (Zeiss) was used to find the cells using transmitted light imaging. Tipped MLCT-C (triangular) AFM cantilevers with a nominal spring constant of 0.005 − 0.02 N/m (Bruker) were employed for the membrane tether-pulling assay. Cantilevers were plasma cleaned for 2 minutes and then incubated in a PBS solution containing 2.5 mg/ml FITC-conjugated Concanavalin A (Sigma, #C7642) for 30 minutes at room temperature. After incubation, they were mounted on the cantilever holder for AFM tether pulling.

The cantilever’s spring constant was calibrated on each measurement day using thermal noise analysis (JPK-SPM software). During the measurement, the cells were maintained at 37°C using a Petri dish heater (JPK Instruments). Cells were measured individually; first, a mitotic cell was selected based on its spherical shape. The cantilever was then approached towards the cell, positioning the tip in the center of the cell. A scan was performed in a 4 × 4 grid pattern (with an edge length of 0.5 *µ*m) over the area located by the eye at the center of the mitotic cell. For each grid point, tether-pulling commenced by lowering the cantilever onto the cell at a speed of 1 *µ*m/s until a set-point force of 1 nN was reached. The cantilever was then held at a constant height for 1 *s* before retracting by 5 *µ*m at the same speed of 1 *µ*m/s.

Throughout the measurements, the force on the cantilever was continuously recorded at a frequency of 10 Hz. This data was subsequently used to measure tether-pulling forces using the ‘step fit’ analysis module of the JPK-SPM Data analysis software, which focused on positive force steps. In Fig. 4c, each tether gave rise to one data point in the statistics. The number of all tethers measured per condition *n*_*T*_ and the number of cells measured *n*_*C*_ are shown in the respective x-axis label.

### G. Solving model equations

To determine the values of membrane tension *γ*_mem_ and filopodia length *L*_fil_, we numerically solved the system of equations defined by Equations (1) and (2) using Mathematica’s NSolve function^54^. We verified that assuming an elastic membrane with elastic reference area *A*_mem,0_ and modulus of *K*_mem_ ≈ 300 mN/m^15^ and using the relation 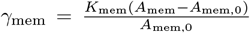 yielded tension estimates deviating by less than 0.1% from those obtained under the inextensibility assumption. This minimal deviation suggests that the inextensibility assumption is valid for our analysis.

## ACKNOWLEDGMENTS

We thank Valentin Ruffine, Michael Schlierf and James Saenz for the good discussions on the project. Furthermore, EFF thanks the Deutsche Forschungsgemeinschaft (DFG, German Research Foundation) for financial support through the Heisenberg program, project number 495224622 (FI 2260/8-1) and the grant FI 2260/5-2. In addition, the authors thank the CMCB Light Microscopy Facility and the PoL Light Microscopy Facility for their excellent support.

## DATA AVAILABILITY STATEMENT

All data generated or analyzed during this study are available from https://doi.org/10.25532/OPARA-757.

## AUTHOR CONTRIBUTIONS

Y.A.P. and E.F.-F. performed the experiments. Y.A.P. and E.F.-F. designed the experiments and performed data analysis. Y.A.P. and E.F.-F. wrote the manuscript.

## SUPPLEMENTARY MATERIAL

**Figure S1.**
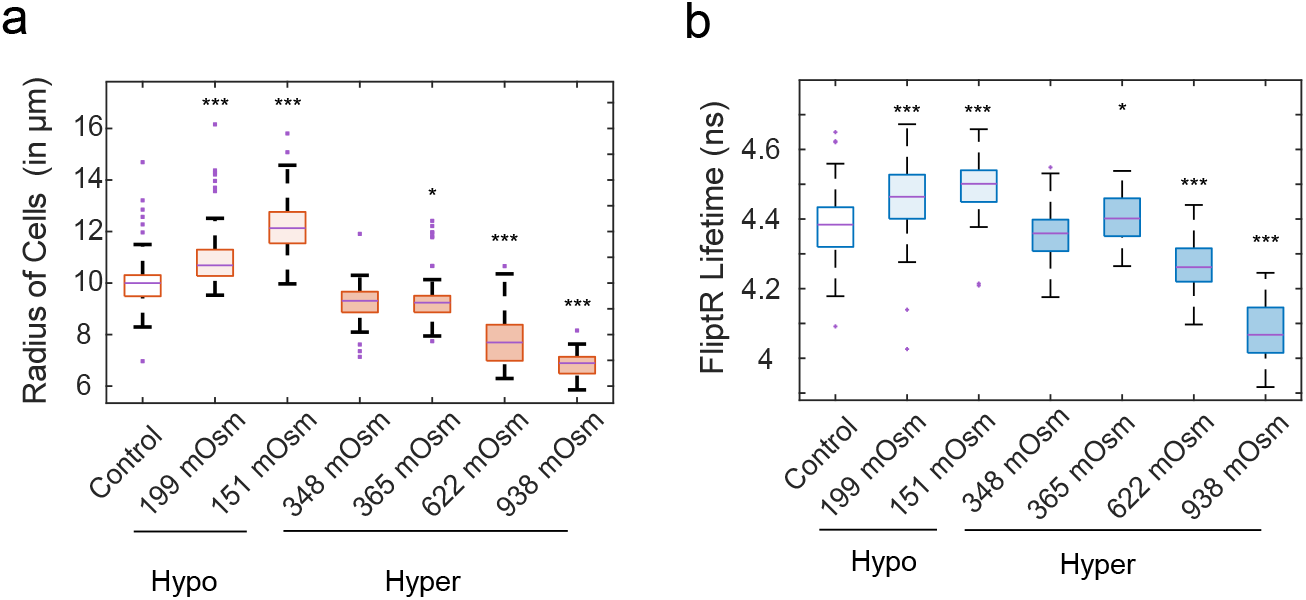
Boxplots of cell radii and FliptR lifetimes following osmotic shock treatment a) The boxplot demonstrates a significant increase in cell radius during hypo-osmotic shock, while a decrease in cell radius is observed during hyper-osmotic shock. Additionally, the change in cell radius is more pronounced as the intensity of the osmotic shock increases in either direction. b) The boxplot depicts the changes in FliptR lifetime upon osmotic shock corresponding to panel a. Cell radii and FliptR lifetimes were averaged in the time interval 500 − 600 s after the shock. We observe that the FliptR lifetime increases with hypo-osmotic shock and decreases with hyper-osmotic shock, indicating a direct dependence between changes in cell size and FliptR lifetime.

**Figure S2.**
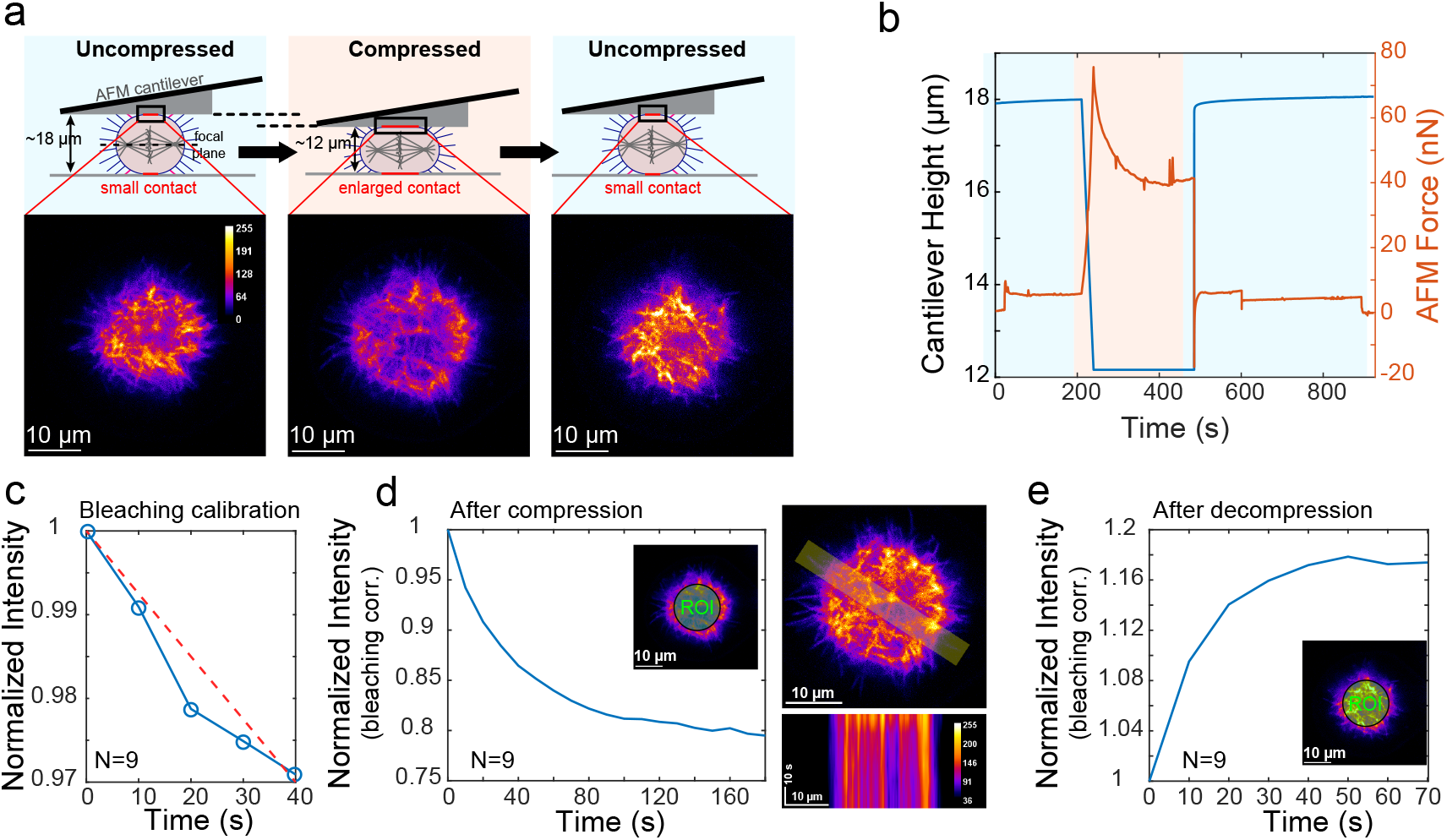
Filopodia intensity declines in newly established contact areas. a) Schematic illustrating the three phases of the experiment. First the cell is kept at shallow confinement (*h* = 18 *µ*m, ‘Uncompressed’) with a small contact area. Then the cantilever is lowered to (*h* = 12 *µ*m, ‘Compressed’) to increase the contact area. In the last phase, the cantilever is retracted back to the original height (*h* = 18 *µ*m, ‘Uncompressed’) releasing cell surface in the focal plane from contact. F-actin intensity (Lifeact-mScarlet) in the upper plane of contact was monitored over time with confocal imaging. The three confocal images show f-actin fluorescence in the plane of contact after 100 s of equilibration time in the respective state. Note that the fluorescent region observed in the confocal images does not entirely correspond to the cell–substrate contact area. Based on shape estimates as previously described^11^, we estimate that in the ‘Uncompressed’ state, only ≲ 10% of the fluorescent region represents actual contact area. This fraction increases to approximately 50% in the ‘Compressed’ state, with some variation depending on cell size. b) Exemplary readout of cantilever height (blue curve) and AFM force (orange curve) during the experiment. c) Intensity evolution during imaging in the contact plane in the first phase of the experiment (‘Uncompressed’) averaged over ten cells. The approximately linear intensity decline was used to estimate the rate of bleaching during imaging. d) F-actin intensity and filopodia abundance visibly reduce in the focal plane upon contact area establishment. Time *t* = 0 s coincides with the time when the cantilever reaches the lowest position. e) F-actin intensity and filopodia abundance visibly increase in the focal plane upon contact area reduction. Time *t* = 0 s coincides with the time point when the cantilever was retracted.

**Caption, Movie 1:** 3D z-stack reconstruction and rotational visualization of a HeLa cell in mitotic arrest highlighting the distribution of filopodia. F-actin was fluorescently labeled with Lifeact-mScarlet to visualize filopodia and retraction fibers. The image stack was acquired using a Zeiss C-Apochromat 40x/1.20 water objective with a spatial resolution of 48.1773 pixels/µm. A total of 137 z-slices were acquired at 0.25 µm intervals using a 561 nm laser, covering a field of view of 42.51 µm × 42.51 µm (2048 × 2048 pixels). The movie begins at the bottom of the cell, visualizing retraction fibers, and rotates 360° around the y-axis to reveal the spatial distribution of filopodia.

**Figure S3.**
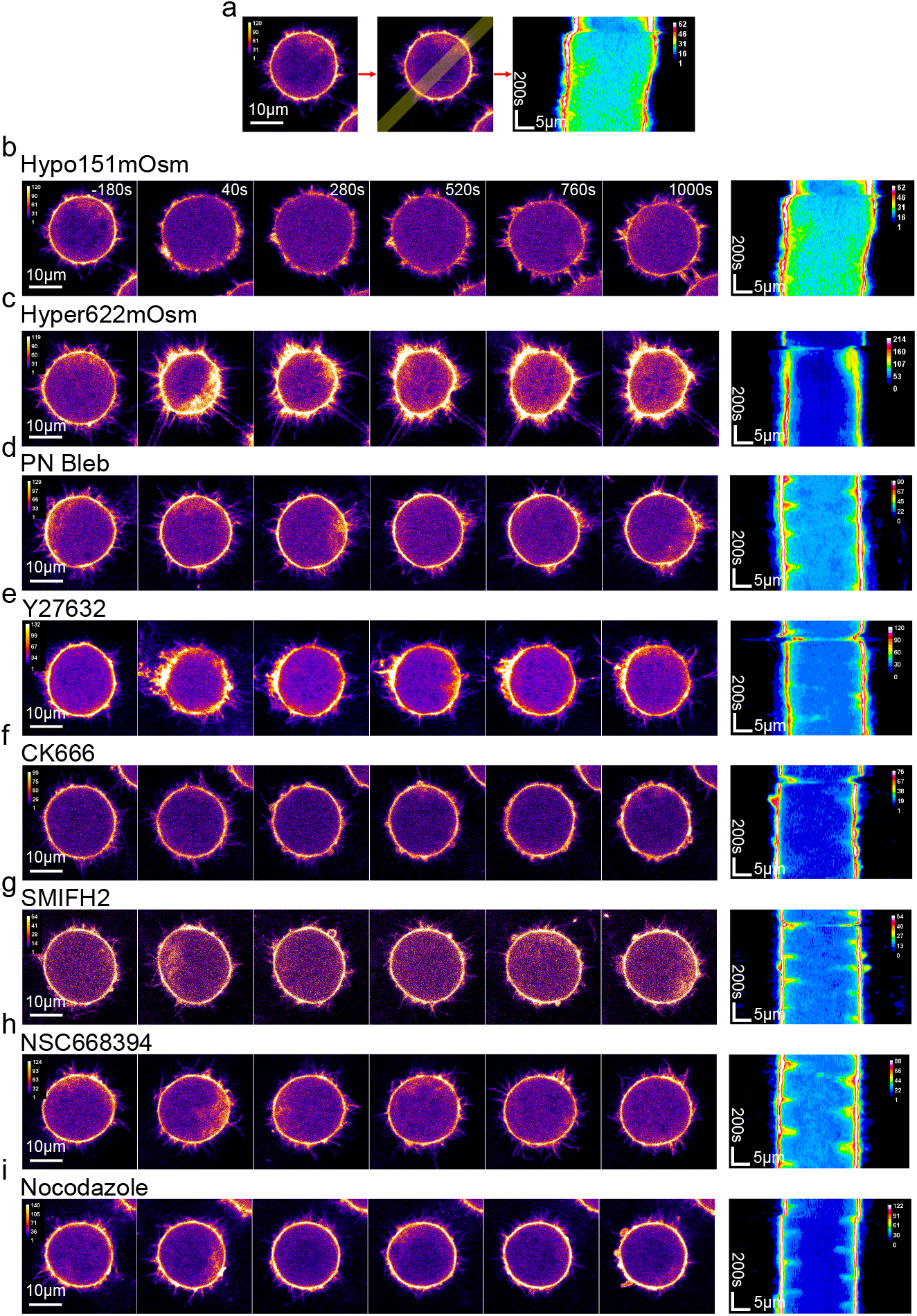
Time dynamics of filopodia membrane reservoirs in HeLa cells during mitotic arrest following osmotic shock and cytoskeletal perturbations. a) A confocal microscopy image of a HeLa cell in mitotic arrest at the equatorial plane displays a representative region that has been selected for kymograph analysis. b-i) A series of confocal micrographs illustrate the visual time dynamics of HeLa cells in mitotic arrest, while the corresponding kymograph demonstrates the time dynamics of filopodia within the selected region following osmotic shock (b & c) and cytoskeletal perturbations (d-i). b) Upon hypo-osmotic shock, we observe an increase in cell size accompanied by a decrease in the number of filopodia, which is evident in both the confocal micrographs and the kymograph. Over time, there is a restoration in the number of filopodia due to active cellular processes. c) In contrast, hyper-osmotic shock results in a decrease in cell size and an increase in the number of filopodia.

**Figure S4.**
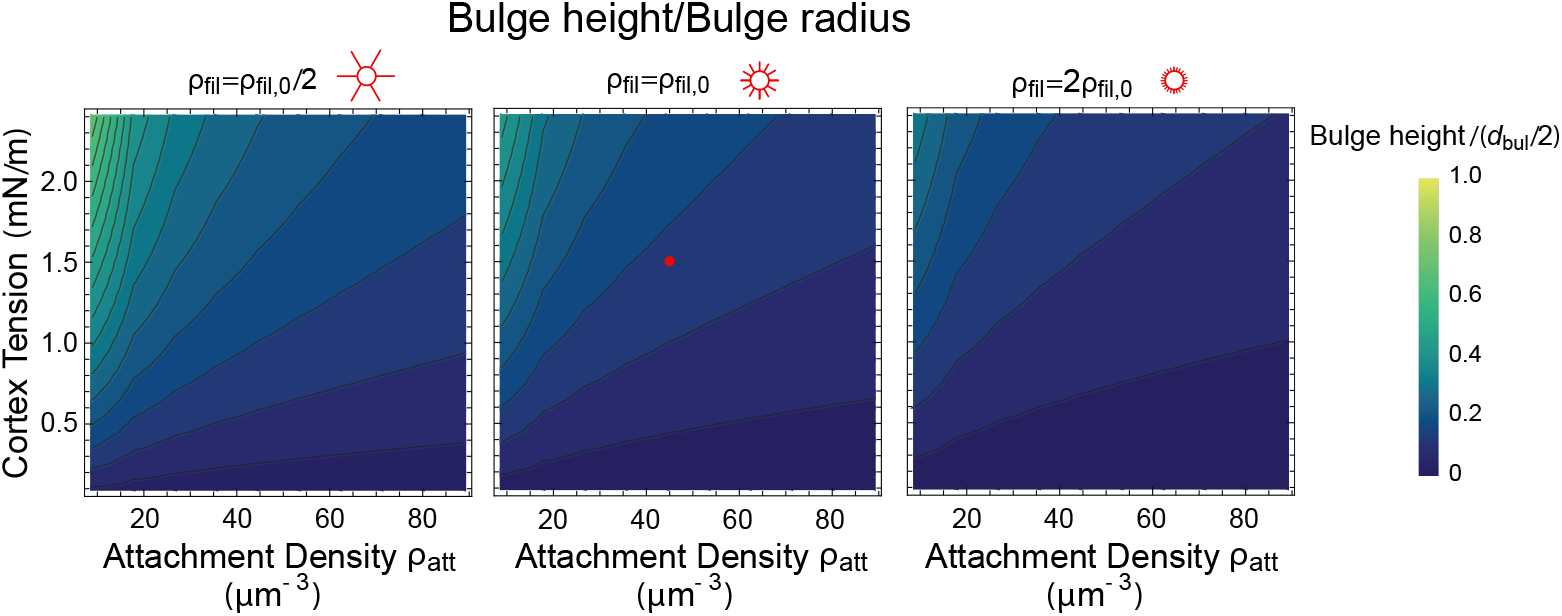
Membrane bulge height normalized by the bulge base radius as predicted by our model in dependence of cortex-membrane attachment *ρ*_*rmatt*_ and cortex tension *γ*_cortex_. Membrane bulges are increasing their depth at higher cortical tensions and lower cortex-membrane attachment. Parameters were chosen as in Fig. 8a,b.

